# GDF3 promotes adipose tissue macrophage-mediated inflammation via altered chromatin accessibility during aging

**DOI:** 10.1101/2024.09.23.614375

**Authors:** In Hwa Jang, Victor Kruglov, Stephanie H. Cholensky, Declan M. Smith, Anna Carey, Suxia Bai, Timothy Nottoli, David A. Bernlohr, Christina D. Camell

## Abstract

Age-related susceptibility to sepsis and endotoxemia is poorly defined, although hyperactivation of the immune system and the expansion of the visceral adipose as an immunological reservoir are underlying features. Macrophages from older organisms exhibit substantial changes, including chronic NLRP3 inflammasome activation, genomic remodeling and a dysfunctional, amplified inflammatory response upon new exposure to pathogen. However, the mechanisms by which old macrophages maintain their inflammatory phenotype during endotoxemia remains elusive. We previously identified *Gdf3*, a TGFβ superfamily cytokine, as a top-regulated gene by age and the NLRP3 inflammasome in adipose tissue macrophages (ATMs). Here, we demonstrate that endotoxemia increases inflammatory (CD11c^+^) ATMs in a *Gdf3-* dependent manner in old mice. Lifelong systemic or myeloid-specific deletion of *Gdf3* leads to reduced endotoxemia-induced inflammation, with decreased CD11c^+^ ATMs and inflammatory cytokines, and protection from hypothermia. Moreover, acute blockade of *Gdf3* using JQ1, a BRD4 inhibitor, phenocopies old mice with lifelong *Gdf3-*deficiency. We show that GDF3 promotes the inflammatory phenotype in ATMs by phosphorylating SMAD2/3. Mechanistically, the differential chromatin landscape of ATMs from old mice with or without myeloid-driven *Gdf3* indicates that GDF3- SMAD2/3 signaling axis shifts the chromatin accessibility of ATMs towards an inflammatory state during aging. Furthermore, pharmaceutical inhibition of SMAD3 with a specific inhibitor of SMAD3 (SIS3) mimics *Gdf3* deletion. SIS3 reduces endotoxemia-mediated inflammation with fewer CD11c^+^ ATMs and less severe hypothermia in old, but not young mice, as well as reduced mortality. In human adipose tissue, age positively correlates with *GDF3* level, while inflammation correlates with pSMAD2/3 level. Overall, these results highlight the importance of GDF3-SMAD2/3 axis in driving inflammation in older organisms and identify this signaling axis as a promising therapeutic target for mitigating endotoxemia-related inflammation in the aged.

## INTRODUCTION

The aging of the immune system and chronic inflammation are key contributing factors to increased hospitalization, mortality and chronic repercussions observed in older individuals with infection-related endotoxemia or sepsis^1–4^. Visceral adipose tissue (VAT) is an organ that exhibits early signs of immune activation during aging and acts as a reservoir for immune cells that play a pivotal role in metabolic or infectious challenges^5–9^. Adipose tissue macrophages (ATMs) are a prevalent immune cell subset that promote chronic inflammation and tissue function decline via the NLRP3 inflammasome activation^10^. However, the mechanism by which aged ATMs maintain their inflammatory phenotype during endotoxemia is unknown. Transcriptome analysis previously identified an age- and NLRP3-dependent upregulation of growth differentiation factor 3 (GDF3) in ATMs, which links GDF3 to age-associated inflammation^11^. Here, we investigate the GDF3-SMAD2/3 signaling axis in endotoxemia-induced inflammation during aging.

GDF3, a cytokine in the TGFβ superfamily, regulates cell fates by modulating the pluripotency of embryonic stem cells^12^. GDF3 has contrasting functions, such as promoting metabolic dysfunction, improving muscle regeneration during injury, and reducing inflammation during sepsis^13–21^, which likely stem from its divergent signaling capabilities^22,23^. Recent studies have correlated GDF3 with disease risk in myocardial infarction, metabolic dysfunction-associated steatohepatitis, obesity, sepsis, and Alzheimer’s disease^13,14,20,24–26^. Like other TGFβ-members, GDF3 signals through type I and type II activin receptors to activate the transcription factors SMAD2 and 3 (SMAD2/3) via phosphorylation^27^. Due to their low binding affinity to DNA, SMAD2/3 elicit divergent transcriptional outcomes, ranging from anti-inflammatory to pro-inflammatory effects depending on the cellular contexts^28–31^. Transcription factors like JUN and FOS family members support increased inflammation during aging, but the contribution of SMADs is unclear^32^.

Our data demonstrate that GDF3 induces an inflammatory phenotype within ATMs, without affecting proliferation and infiltration, from old mice during endotoxemia. Lifelong deletion of *Gdf3*, at a whole-body or myeloid-specific level, reduces inflammatory macrophages, inflammatory cytokines, and hypothermia in response to endotoxemia. Mechanistically, GDF3 increases inflammatory ATMs by phosphorylating SMAD2/3. GDF3 induces chromatin accessibility alterations in ATMs from old mice, which are reversed with myeloid-specific *Gdf3* deletion. Two distinct pharmaceutical approaches targeting the GDF3-SMAD2/3 axis alleviate endotoxemia-induced inflammation in old mice. Moreover, SMAD3-blockade protects against endotoxemia-induced lethality in old mice. In human VAT, we find that age positively correlates with *GDF3* level, and inflammatory cytokine expression correlates with pSMAD2/3 level. These data collectively suggest that GDF3-SMAD2/3 axis plays a critical role in shaping the inflammatory phenotype of ATMs during endotoxemia and aging, and targeting this pathway mitigates endotoxemia-associated inflammation in old mice.

## RESULTS

### Endotoxemia is associated with an increase in pro-inflammatory adipose tissue macrophages in old mice

Older individuals exhibit increased lethality, cytokine storm and impaired metabolic responses to sepsis and endotoxemia, partly mediated by immune cell crosstalk within VAT^6,33,34^. To address whether the increase in inflammation is associated with a change in macrophage phenotype, we investigated the phenotype of ATMs at 4 hours after administering PBS or 0.1mg/kg LPS via intraperitoneal (i.p.) injection to young or old wildtype (WT) mice (Figure 1A, left). Consistent with previous studies^6^, old mice failed to maintain core body temperature (BT) with this low-dose LPS challenge (Figure 1A, right). We initially examined the phenotype of ATMs using multi-parameter flow cytometry (representative gating strategy in extended data), with CD11c as a marker for inflammatory phenotypes. After LPS injection, the frequency of CD11c^+^ ATMs expanded in old mice compared to PBS injection, while it remained unchanged in young mice (Figure 1B, S1A and S1B). To further characterize CD11c^+^ ATMs, we analyzed multiple markers generally used to define macrophage phenotypes: CD11c, CD9, MHCII and CD206. The tSNE plot shows that ATMs are a heterogeneous population with varying combination of markers (Figure 1C). Notably, CD11c^+^ and CD206^+^ ATMs, commonly categorized as pro- and anti-inflammatory ATMs, represent distinct populations. MHCII and CD9 were expressed on CD11c^+^ ATMs, further supporting CD11c as a marker for inflammatory ATMs^35^. These data indicate that low-dose LPS exacerbates the imbalance of a heterogenous population of inflammatory CD11c^+^ ATMs in an age-dependent manner. To further investigate the phenotype of VAT immune cells, we utilized single-cell RNA sequencing data set (GSE274935) containing VAT CD45^+^ immune cells sorted from young and old mice that were challenged with low-dose LPS. Young and old VAT immune cells were analyzed together and a total of 23 clusters were identified including 4 clusters of ATMs (ATM1, 2, 3, and 4; Figure 1D). The proportions of each ATM cluster were altered during aging. The frequency of ATM1 cluster decreased from 71% to 51%, while the ATM2 cluster increased from 20% to 32% and the ATM3 cluster increased from 5% to 14% with age (Figure 1E). The ATM2 and ATM3 cluster exhibited higher expressions of *Itgax* (CD11c), *Cd9*, MHCII-associated genes (*H2-Ab*), while showing lower expression of *Mrc1* (CD206) and *Lyve1* (Figure 1F). These characteristics of ATM2 and ATM3 cluster are comparable to those of inflammatory CD11c^+^ ATMs identified by flow cytometry, as shown in Figure 1C. Moreover, these clusters exhibit higher expression of *Gdf3* compared to other ATM clusters, suggesting that inflammatory ATMs are the primary source of *Gdf3*.

**Figure 1.**
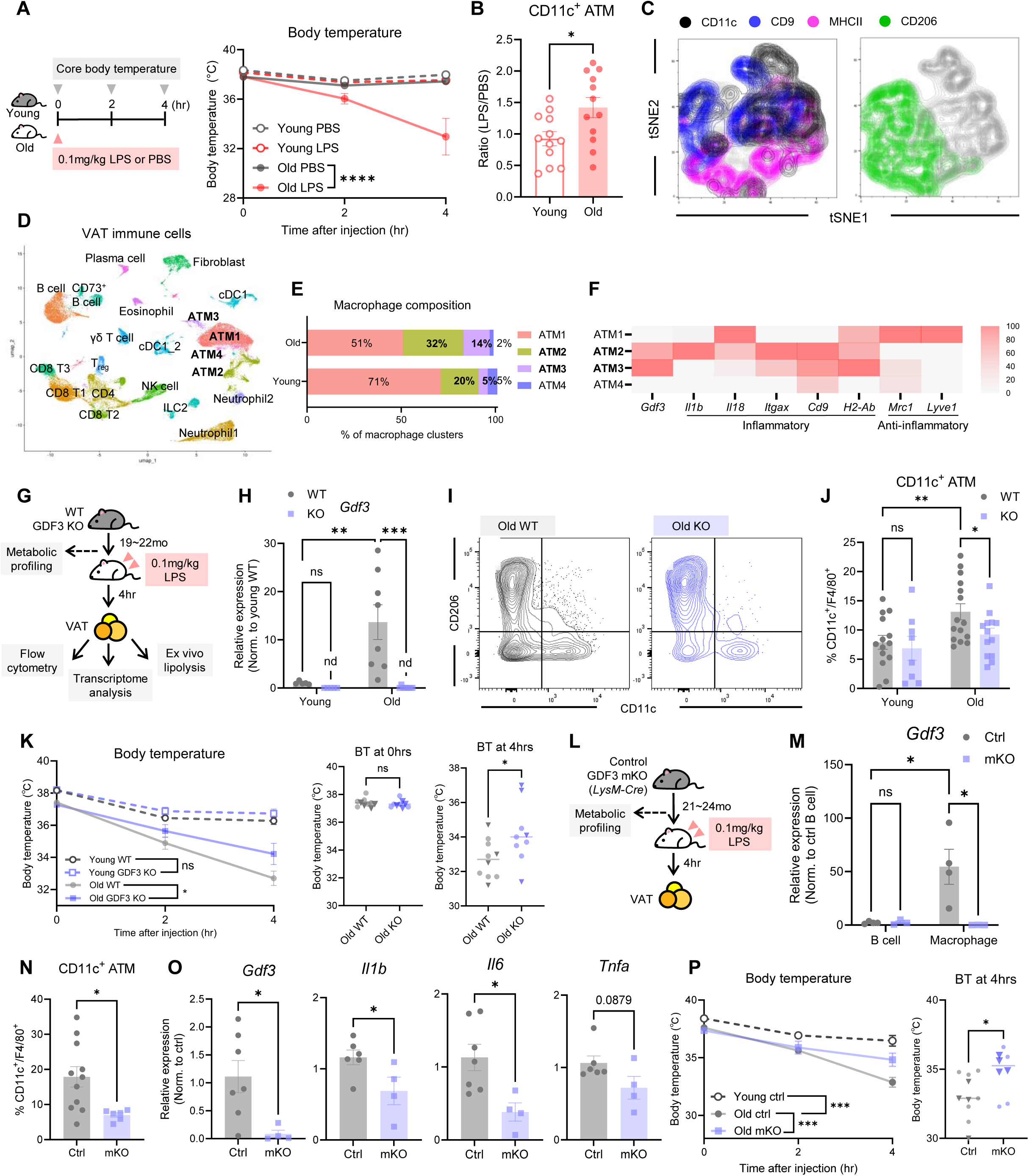
*Gdf3* promotes endotoxemia-induced inflammation by regulating inflammatory ATMs during aging. (A) Schematic of experimental design (left). Core BT of young (3-month-old) or old (22-month-old) female WT mice after i.p. injection of 0.1mg/kg LPS or PBS (right). n=3 per group. (B) Ratio of CD11c^+^ ATMs (% of CD11c^+^ ATMs from LPS-injected group normalized to PBS-injected group). (C) tSNE plot of compiled F4/80^+^ ATMs in VAT from 24-month-old female WT mice (n=5) injected with 0.1mg/kg LPS. 1000 iterations, 30 perplexities. (D) UMAP of VAT immune cells (CD45^+^ cells) from young (3-month-old) or old (21-month-old) female WT mice injected with 0.1mg/kg LPS showing 23 clusters. (E) Proportion of each ATM clusters (% of all young or old ATMs) (F) Expressions of genes associated with macrophage phenotype to compare ATM clusters (ATM1, 2, 3, 4). (G) Schematic of experimental design. (H-K) Young (3- to 4-month-old) or old (19- to 22-month-old) female WT or GDF3 KO mice injected with 0.1mg/kg LPS. (H) Gene expression of *Gdf3* in VAT. Young WT (YW), n=5; young KO (YK), n=5; old WT (OW), n=8; old KO (OK), n=8. (I) Representative gating strategy for CD11c^+^ and CD206^+^ ATMs in VAT from OW (left) or OK (right) mouse. (J) Frequency of CD11c^+^ ATMs. YW, n=14; YK, n=8; OW, n=15; OK, n=14. (K) Mean core BT after LPS injection. YW, n=14; YK, n=8; OW, n=8; OK, n=8. Circle and triangle symbols represent two independent experiments. (L) Schematic of experimental design. (M) Gene expression of *Gdf3* in positively selected B cells from spleen and peritoneal macrophages. Ctrl, n=4; mKO, n=3. (N-P) Young (4-month-old) or old (21- to 24-month-old) female control or mKO mice were injected with 0.1mg/kg LPS. (N) Frequency of CD11c^+^ ATMs and (O) inflammatory cytokine gene expression in VAT from old control (n=6∼11) or mKO mice (n=4∼6). (P) Mean core BT after LPS injection. Young control, n=8; old control, n=11; old mKO, n=8. Circle and triangle symbols represent two independent experiments. All data are presented as means ± SEM. *p<0.05, **p<0.01, ***p<0.001, ****p<0.0001. All *in vivo* experiments were repeated independently at least twice. ns, not significant. nd, non-detected. See also Figure S1 and S2.

### Old *Gdf3*-deficient mice are protected from endotoxemia-induced hypothermia and the increase in pro-inflammatory macrophages

GDF3 is implicated in age-related adipose tissue inflammation and affects severity or risk for multiple diseases^11,14,20,21,24,26,36^. We next asked whether lifelong deficiency of *Gdf3* would alter the endotoxemia-related inflammatory state of macrophages in VAT from old mice (Figure 1G). To address this question, we generated mice with whole-body deletion of *Gdf3.* We first validated deletion of *Gdf3* in VAT from young (3- to 4-month-old) or old (19- to 22-month-old) WT or *Gdf3* whole-body knockout (KO) mice. The result showed a significant age-related increase in *Gdf3* expression that was ablated in old KO mice (Figure 1H). To investigate the impact of lifelong *Gdf3* deletion on metabolic function prior to endotoxic shock, we monitored various metabolic parameters, including body weights, body composition, insulin sensitivity and glucose sensitivity. However, we only observed an age effect without a genotype effect (Figure S1C-F). Genes related to fatty acid oxidation or synthesis also showed no differences in adipocytes isolated from VAT (Figure S1G). Taken together, lifelong *Gdf3*-deficiency has no impact on metabolic dysfunction.

To evaluate the impact of *Gdf3*-deficiency on immune cell activation and ATMs’ inflammatory phenotype during endotoxemia, we quantified the frequency of lymphocytes and CD11c^+^ or CD206^+^ ATMs from the VAT. When young and old WT mice were compared, T cells, B cells and myeloid cells, including ATMs, exhibited the expected age-related changes in both frequency and cells per gram of tissue (Figure S1H, cellularity data not shown). However, no genotype effect was observed in this overall immune composition between old WT and KO mice (Figure S1H). Next, we examined the inflammatory profile of ATMs (Figure 1I). Notably, deletion of *Gdf3* resulted in a decreased frequency of CD11c^+^ ATMs, while the frequency of CD206^+^ ATMs remained unaffected (Figure 1J and S1I).

In the spleen, there were no differences in immune composition identified between old WT and KO mice, indicating that the effect of *Gdf3* is primarily within VAT (Figure S1J). Impaired lipolysis upon aging leads to dysregulated immune response^6^. As GDF3 can limit adipocyte lipolysis, we measured free fatty acids released from VAT explants^11,14,15^. VAT explants from old KO mice exhibited improved stimulated-lipolytic capacity compared to their old counterparts (Figure S1K). Next, we assessed the severity of endotoxemia by monitoring core BT at 0, 2, and 4 hours post-LPS injection. While there were no differences in core BT in young WT and KO mice, old WT mice exhibited a greater reduction in BT compared to old KO mice (Figure 1K). Taken together, these results suggest that GDF3 mediates endotoxemia-induced inflammation in old mice by regulating the inflammatory phenotype of ATMs, lipolysis and hypothermia.

To address how GDF3 induces the expansion of CD11c^+^ macrophages, we investigated whether this increase occurs via proliferation or infiltration. As GDF3 promotes proliferation of various cell types^19,24^, we first hypothesized that the expansion of LPS-induced CD11c^+^ ATMs was dependent on GDF3-regulated proliferation. To perform an *in vivo* cell proliferation study, young (4-month-old) and old (24-month-old) WT mice were i.p. injected with EdU (5-ethynyl- 2’deoxyuridine, 25 mg/kg) for 4 consecutive days, followed by PBS or 0.1mg/kg LPS injection (Figure S1L)^37^. Cell proliferation was assessed by quantifying incorporation of EdU into DNA by flow cytometry (Figure S1M). The frequency of EdU^+^CD11c^+^ ATMs was elevated 1.7-fold with LPS injection in old mice, while it remained unchanged in young mice (Figure S1N). To test whether *Gdf3*-deficiency would reduce the proliferation of CD11c^+^ ATMs, WT and KO mice were injected with EdU (Figure S1O, left). Although EdU^+^CD11c^+^ ATMs were elevated with aging, no differences between old WT and KO mice were observed (Figure S1O, right). Monocytes can infiltrate into VAT and differentiate into inflammatory macrophages^38^. To investigate whether GDF3 influences infiltration of monocytes, we used the CCR2 as a marker for infiltrating monocytes^39^. While the frequency of CCR2^+^ ATMs increased with age, we found no differences between old WT and KO mice (Figure S1P). Collectively, our data demonstrate that GDF3 induces inflammatory phenotype of ATMs independently of proliferation or infiltration, despite both being crucial for maintaining CD11c^+^ ATMs during endotoxemia in old mice.

### *Gdf3* is not required for LPS-induced lethality in young mice, but is regulated with age in a B cell-dependent manner

We next asked whether the protective function of *Gdf3* is exclusive to old mice. Young WT and KO mice (aged 4- to 6- month-old) were injected with a lethal dose of LPS (18mg/kg) and monitored for 6 hours post-LPS administration (Figure S2A). Core BT, immune cell composition, lymphocyte activation, and inflammatory gene expression remained unaltered in young mice regardless of the presence of *Gdf3*, indicating that *Gdf3* does not play a role in severe inflammatory responses in young mice (Figure S2B-F). These findings suggest that the beneficial effect from *Gdf3* deletion against severe inflammatory responses is specific to old mice.

To broadly address age-related alterations in expression of *Gdf3* and its family members, we first compared the gene expression of selected *Tgfb*-associated genes^23^. Consistent with our observations in Figure 1H, *Gdf3* increased with age in VAT. In line with previous reports, *Gdf15* was elevated, but not statistically significant^40^. There were no differences detected in other genes from TGFβ family (Figure S2G). Moreover, *Gdf3* expression was elevated in multiple organs of old mice, with the highest expression observed in VAT, liver and thymus compared to young mice (Figure S2H). Lifelong B cell-deficiency improves aged adipose tissue microenvironment by reducing inflammatory ATMs^6,41^. Notably, old B^null^ mice exhibited reduced *Gdf3* and *Tgfb1,* but not *Gdf15,* expression compared to old WT mice (Figure S2I). These findings suggest that *Gdf3* is highly expressed in VAT and increases in a B cell-dependent manner during aging.

### Myeloid-driven *Gdf3* is required for endotoxemia-induced cytokine storm, inflammatory macrophages and hypothermia

As *Gdf3* may be expressed in multiple cell types, we wanted to determine the effects of deleting *Gdf3* in a myeloid cell-specific manner. *Gdf3*-floxed mice were crossed with *LysM-Cre* line to generate myeloid cell-specific knockout (mKO) mice and control littermates. We first measured *Gdf3* expression to confirm the *Cre*-mediated deletion. Macrophages from peritoneum exhibited a reduction of *Gdf3* expression in mKO group. In contrast, B cells showed comparable levels of *Gdf3* expression between control and mKO mice, validating the myeloid-specific deletion of *Gdf3* (Figure 1L and 1M). Although mKO mice showed no differences in body weights, they showed improved glucose sensitivity at 20- to 23- months of age when compared to control mice (Fig S2J-M).

Next, we predicted that myeloid-driven *Gdf3* is required for the inflammatory phenotype of macrophages during endotoxemia. At 21- to 24-months of age, control and mKO mice were given 0.1mg/kg LPS (Figure 1L). ATMs from the old mKO group showed a significant reduction in the frequency of CD11c^+^ ATMs (Figure 1N). There was also a significant decrease in CD9^+^ ATMs, a slight increase in CD206^+^ and no change in CCR2^+^ ATMs (Figure S2N-P). The decreased frequency in CD11c^+^ macrophages was also seen in the subcutaneous adipose tissue, indicating the conserved function of *Gdf3* across the white adipose depots (Figure S2Q). *Gdf3* expression was significantly reduced in VAT from mKO mice, indicating that myeloid cells are the main contributors of *Gdf3* to the tissue (Figure 1O). To evaluate the impact of reduced CD11c^+^ ATMs, we measured inflammatory cytokine gene expression (*Il1b, Il6, Tnfa*). These genes were decreased in VAT from old mKO mice compared to control mice (Figure 1O). Moreover, stimulated-lipolytic capacity of VAT explants was improved with myeloid-specific deletion of *Gdf3* (Figure S2R).

In aged VAT, B cells regulate adipose tissue homeostasis. They contribute to elevated inflammation by modulating other immune cells, including ATMs, and by inhibiting stimulated lipolysis^6,42^. We asked if altered ATMs would affect B cells. The overall frequency of B cells showed no differences between old control and mKO mice (data not shown). To further characterize B cells, we used IgD, IgM, programmed cell death ligand 1 (PDL1) and CD73 as markers for dysfunctional B cell in aged VAT (Figure S2T)^6^. Markedly, both IgM^+^CD73^+^ and PDL1^+^ B cells were decreased in old mKO mice (Figure S2U and S2V). Collectively, this suggests that *Gdf3* can modulate dysfunctional B cells that accumulate with age in VAT.

During endotoxemia old control mice developed hypothermia with 13% BT loss, whereas old mKO mice had significantly increased BT at 4 hours post-endotoxemia, showing 7% BT loss (Figure 1P). This was also observed in male mKO mice that displayed improved glucose sensitivity and fewer CD11c^+^ ATMs during endotoxemia, demonstrating a conserved function of *Gdf3* across sexes (Figure S2L, M, S, W). Taken together, our findings suggest that lifelong myeloid-specific deletion of *Gdf3* results in improved glucose sensitivity, reduced CD11c^+^ ATMs, improved lipolytic capacity, decreased inflammatory gene expression and protection from BT loss post-LPS challenge.

### BRD4-controlled *Gdf3* regulates endotoxemia-induced inflammation

Next, we wanted to examine if the acute depletion of *Gdf3* could alleviate inflammation. Bromodomain and extra terminal domain 4 (BRD4), a transcriptional and epigenetic regulator, binds to promoter and enhancer regions of *Gdf3,* facilitating transcription of *Gdf3* in obesity^43^. To first assess whether the expression of *Gdf3* is regulated by BRD4 in older organisms, we utilized JQ1, a small molecular inhibitor of BRD4^44^. At 23- to 24-months of age, mice were i.p. injected with JQ1 or vehicle for 6 consecutive days, followed by 0.1mg/kg LPS injection (Figure 2A). Acute administration of JQ1 reduced *Gdf3* in the aged VAT without affecting body weights (Figure 2B and S3A). We next tested if acute depletion of *Gdf3* would have similar effects on macrophage phenotypes to those observed with lifelong deletion of *Gdf3*. Although the composition and cellularity of overall immune cells in VAT were not affected by JQ1 (Figure S3B and S3C), the composition of ATMs was altered. Specifically, the frequency of CD11c^+^ ATMs decreased, while the frequency of CD206^+^ ATMs increased (Figure 2C). The frequency of CCR2^+^ ATMs remained unaffected (Figure 2D). Gene expression of inflammatory cytokines *(Il18, Il1b, Casp1, Tnfa*) was reduced in VAT (Figure 2E). In line with the observed changes in B cell phenotypes resulting from lifelong myeloid-specific *Gdf3*-deficiency, we found a reduction in PDL1^+^ B cells following JQ1 injection (Figure S3D). However, the frequency of CD73^+^ B cells was not affected (Figure S3E). Moreover, JQ1- injected old mice were protected from hypothermia compared to vehicle-injected mice (Figure 2F).

**Figure 2.**
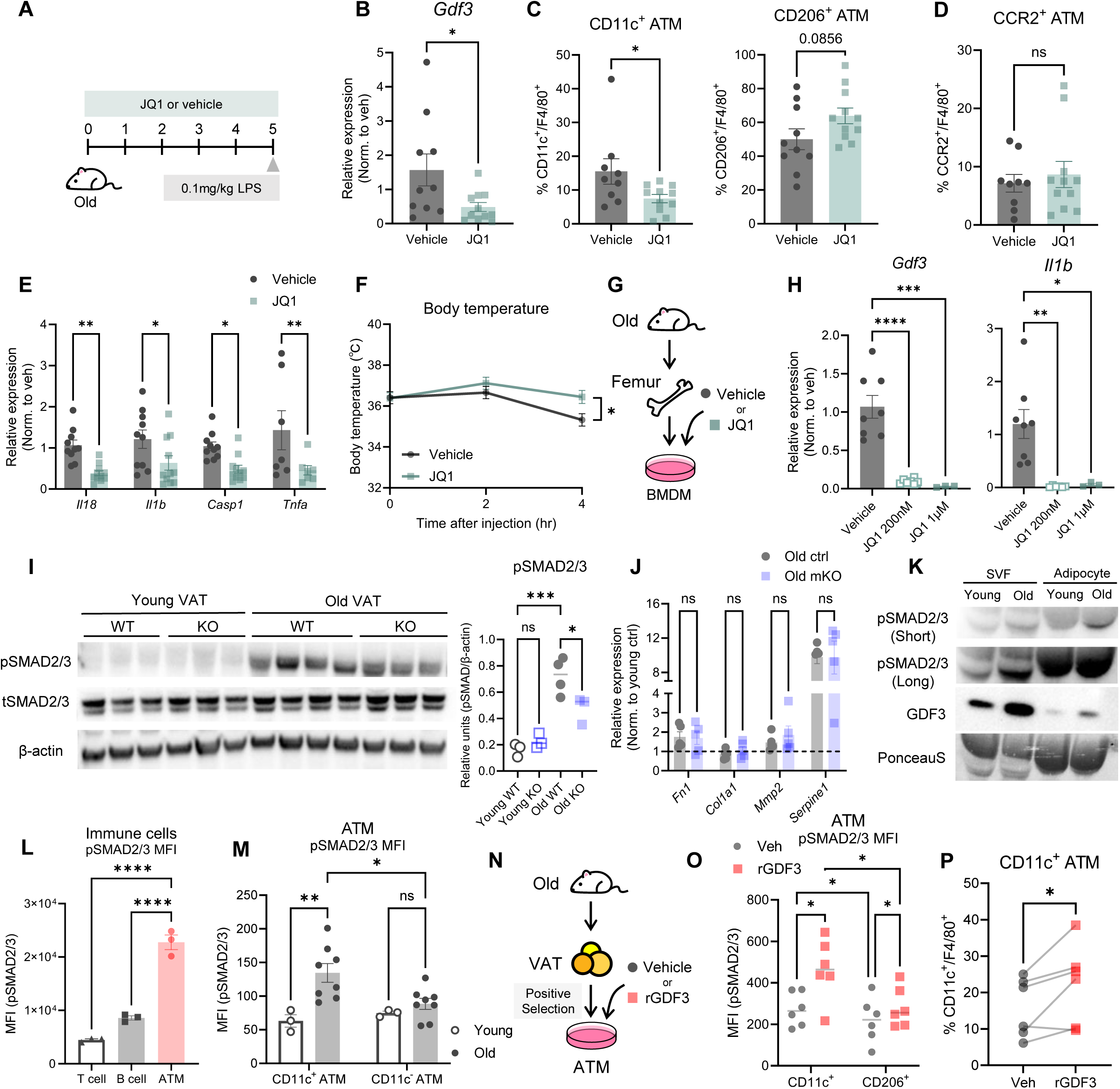
BRD4-regulated *Gdf3* induces inflammatory phenotype of ATM through SMAD2/3 signaling axis in the aged. (A-F) Old (23- to 24-month-old) female WT mice were i.p. injected with JQ1 or vehicle and LPS (0.1mg/kg). Vehicle-injected, n=10; JQ1-injected, n=11. (A) Schematic of experimental design. (B) *Gdf3* expression in VAT. (C) Frequency of CD11c^+^, CD206^+^, and (D) CCR2^+^ ATMs in VAT. (E) Inflammatory gene expression in VAT. (F) Mean core BT after LPS injection. (G-H) BMDMs isolated from 24-month female WT mice were treated with vehicle or JQ1 (200nM or 1µM). (G) Schematic of experimental design. (H) *Gdf3* and *Il1b* expression in BMDMs. (I) Phospho-SMAD2/3 (pSMAD2/3), total-SMAD2/3 (tSMAD2/3), and β-actin by western blot of whole VAT from young (3-month) or old (24-month) female WT and GDF3 KO mice injected with 0.1mg/kg LPS. (J) VAT gene expression involved in fibrosis from old control (n=5) and old mKO (n=5) male mice injected with 0.1mg/kg LPS. Dotted line, young control. (K) pSMAD2/3, GDF3 and β-actin by western blot, and PonceauS staining of SVF or adipocytes from young or old female WT mice injected with 0.1mg/kg LPS. (L) Mean fluorescence intensity (MFI) of pSMAD2/3 in T cells, B cells and ATMs in VAT from 24-month female WT mice (n=3). (M) pSMAD2/3 MFI in CD11c^+^ or CD11c^-^ ATMs from young (4-month-old) and old (23-month-old) female WT mice. Young, n=3; old, n=8. (N-P) Positively selected ATMs from VAT of 24-month female WT mice were treated with vehicle or rGDF3 (20ng/mL) *in vitro*. (N) Schematic of experimental design. (O) pSMAD2/3 MFI in CD11c^+^ or CD206^+^ ATMs, and (P) frequency of CD11c^+^ ATMs treated with vehicle or rGDF3. All data are presented as means ± SEM. *p<0.05, **p<0.01, ***p<0.001, ****p<0.0001. All *in vivo* and *in vitro* experiments were repeated independently at least twice. ns, not significant. See also Figure S3.

To directly test the impact of JQ1 on macrophages, we utilized bone marrow-derived macrophages (BMDMs). BMDMs collected from old mice were treated with an increasing dose of JQ1 or vehicle for 4 hours (Figure 2G). Consistent with *in vivo* JQ1 administration, JQ1 reduced *Gdf3* expression at both doses *in vitro* (Figure 2H, left). We also identified a reduction in the inflammatory genes, *Il1b, Casp1* and *Tnfa* (Figure 2H and S3F). Collectively, data suggest that acute depletion of *Gdf3* via BRD4 inhibition phenocopies lifelong deletion of *Gdf3* in old mice.

### GDF3 induces phosphorylation of SMAD2/3 to promote inflammatory phenotype of ATMs during aging

Previous studies have demonstrated that GDF3 can activate SMAD2/3 signaling via phosphorylation in primary myoblasts and adipocytes^19,43^. We wanted to test if the age-related increase in pSMAD2/3 requires GDF3. There were no differences when comparing pSMAD2/3 in VAT from young WT and KO mice. However, old KO mice showed a reduced pSMAD2/3 levels when compared to old WT mice (Figure 2I). SMAD2/3 signaling promotes fibrosis in multiple tissues including adipose tissue and kidney^41,45^. However, we did not detect differences in the expression levels of fibrotic genes downstream of SMAD2/3 (*Fn1, Col1a1, Mmp2, Serpine1*) between old control and mKO mice in VAT (Figure 2J). Considering the cellular heterogeneity of VAT, we sought to identify which cell types express pSMAD2/3 with aging. The whole VAT from young and old WT mice was separated into adipocyte fraction and stromal vascular fraction (SVF), which includes immune cells. Interestingly, both adipocyte fraction and SVF showed increase in pSMAD2/3 and GDF3 with age (Figure 2K). Next, to determine the immune cells exhibiting elevated expression of pSMAD2/3, we employed phospho-flow cytometry. The expression level of pSMAD2/3, represented by mean fluorescence intensity (MFI), was the highest in ATMs compared to B cells and T cells from aged VAT (Figure 2L). Moreover, we observed an age-dependent elevation in pSMAD2/3 levels specifically within CD11c^+^ ATMs, with no significant changes in CD11c^-^ ATMs (Figure 2M).

To directly examine if GDF3 can induce phosphorylation of SMAD2/3 in macrophages, we used an *in vitro* system with BMDMs and ATMs (Figure 2N and S3G). First, BMDMs from young and old mice were treated with vehicle or recombinant GDF3 (rGDF3, 20ng/mL). At basal levels, pSMAD2/3 was higher in BMDMs from old mice compared to those from young mice (Figure S3H). With rGDF3 treatment, pSMAD2/3 elevated in BMDMs from old mice, but not young (Figure S3H). rGDF3 treatment also reduced the frequency of CD206^+^ BMDMs, and increased CD9 MFI (Figure S3I and S3J). For a more physiologically relevant *in vitro* model, we isolated ATMs using positive selection from SVF of old mice and treated them with vehicle or rGDF3 (Figure 2N). Similar to the increase that is induced by natural aging, CD11c^+^ ATMs exhibited higher basal level of pSMAD2/3 than CD206^+^ ATMs *in vitro* (Figure 2O). Both the CD206^+^ and CD11c^+^ ATMs showed increased pSMAD2/3 following rGDF3 treatment, but CD11c^+^ ATMs exhibited a more robust response to rGDF3 with higher pSMAD2/3 MFI (Figure 2O). Additionally, rGDF3 treatment led to an increase in the frequency of CD11c^+^ ATMs (Figure 2P). Taken together, our data demonstrate that GDF3 increases pSMAD2/3 and has no effect on fibrotic genes, but instead promotes the inflammatory phenotype of macrophages from old mice.

### *Gdf3* regulates chromatin accessibility through SMAD-regulated transcription

SMAD2/3 transcription factors (TFs) have low DNA binding affinity for a specific consensus sequence, allowing them to elicit anti- or pro-inflammatory responses depending on the cellular context^28,30,31,46^. Genes associated with pro-inflammatory processes in immune cells, including myeloid cells, exhibit increased chromatin accessibility with age ^32,47^. We hypothesized that this could affect the transcriptional activity of SMAD2/3. To assess chromatin accessibility, ATMs isolated via fluorescence-activated cell sorting (FACS) from young control, old control and old mKO mice were profiled using assay for transposase-accessible chromatin with sequencing (ATAC-seq; Figure 3A, S4A and S4B). The quality of data was validated through the enrichment of ATAC-seq reads around transcription start sites (TSS) and the genomic distribution pattern of identified peaks that map to promoter and intergenic regions (Figure S4C and S4D). ATAC- differential peak analysis of young and old control ATMs identified 4681 genes with increased accessibility and 1013 genes with decreased accessibility, revealing markedly altered chromatin accessibility with age (Figure 3B). Hierarchy clustering analysis also demonstrated distinctive patterns of chromatin accessibility of young and old ATMs (Figure 3C). To investigate which genes are regulated by *Gdf3*, we next compared ATMs from old control and old mKO mice. Interestingly, the chromatin accessibility of 3248 genes was reduced with *Gdf3* deletion in old ATMs (Figure 3D). Moreover, clustering analysis revealed distinct patterns between old mKO and control ATMs (Figure 3E). Out of 3248 closing peaks with *Gdf3* deletion, 50% (n=1627) overlapped with opening peaks with aging (Figure S4E). Utilizing databases such as KEGG and Gene Ontology (GO), we aimed to identify enriched pathways associated with genes showing differential accessibility with aging or genotype^48,49^. KEGG analysis identified enrichment of ECM-receptor interaction, Th17 cell differentiation, AGE-RAGE signaling complication, Rap1 signaling, PI3K-Akt signaling, cytokine-cytokine receptor interaction, and calcium signaling pathways and others among genes showing increased accessibility with aging (Figure 3F). Remarkably, of these enriched pathways with aging, several overlapped with enriched pathways with genes exhibiting decreased accessibility in mKO ATMs (Figure 3G). These included ECM-receptor interaction and Rap1 signaling pathway that would be expected to be enriched with aging based on previous findings^50–52^. When examining specific cytokine receptor-relevant genes, we observed significant increases in chromatin accessibility around TSS of inflammatory cytokine receptors, including *Il1r1* and *Il18r1*, in ATMs from older mice (Figure 3H). Conversely, the same region became less accessible without *Gdf3* in old ATMs.

**Figure 3.**
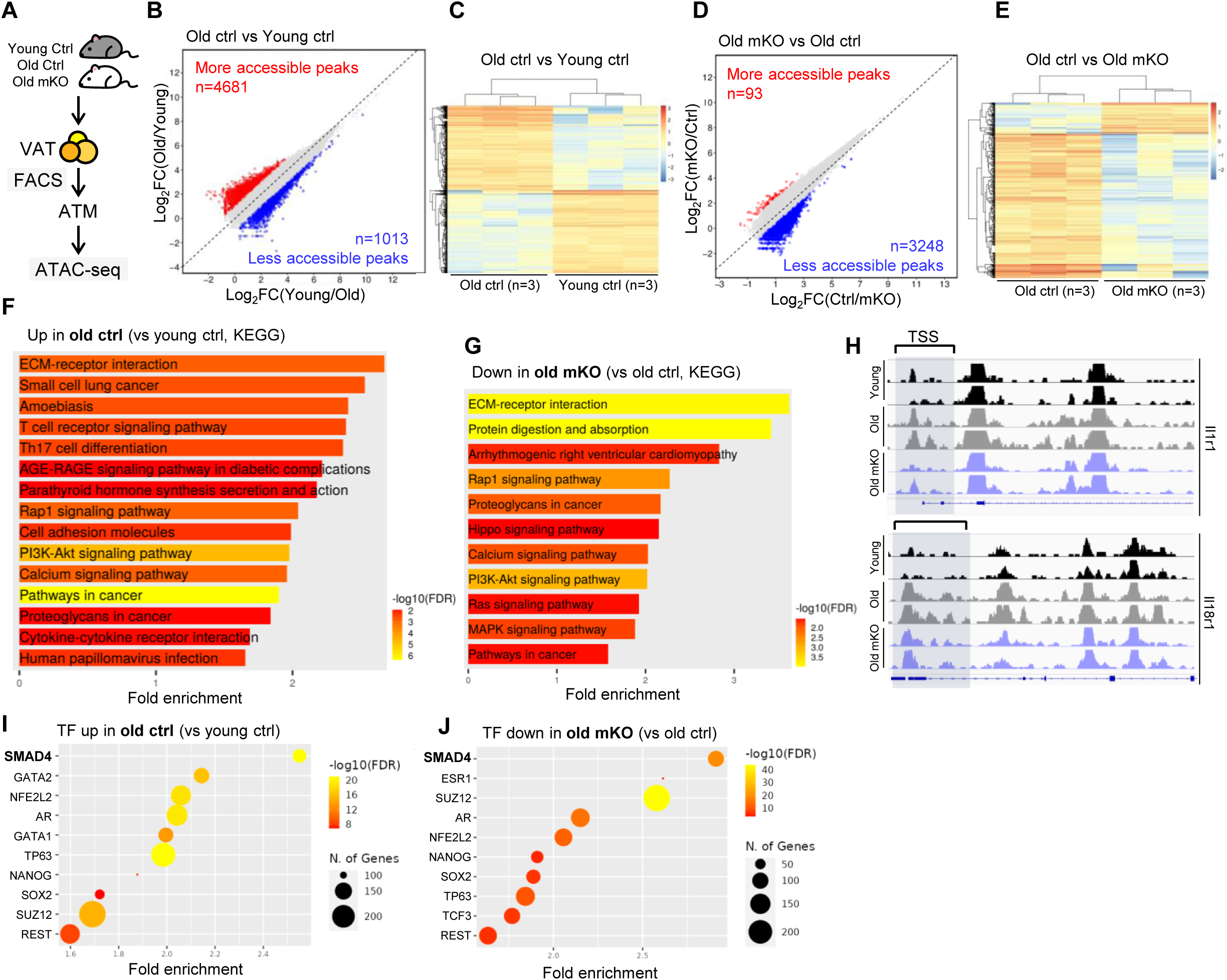
*Gdf3* promotes SMAD-regulated genes via altered chromatin accessibility in ATMs. (A-J) Young (4-month-old) or old (21- to 24-month-old) female control or mKO mice were injected with 0.1mg/kg LPS. (A) Schematic of experimental design. (B-C) Differentially accessible peaks between old and young control ATMs represented by (B) log_2_FC plot and (C) hierarchy clustering heatmap. For log_2_FC plot, differentially opening (closing) peaks are shown in red (blue). (D-E) Differentially accessible peaks between old mKO and old control ATMs represented by (D) log_2_FC plot and (E) hierarchy clustering heatmap. (F-G) KEGG pathway analysis of (F) opening peaks with aging (old vs young control) and (G) closing peaks with *Gdf3* deletion (old mKO vs old control). (H) IGV genome browser tracks that show chromatin accessibility of *Il1r1* and *Il18r1* in ATMs. TSS highlighted with gray boxes. (I-J) Prediction of TF binding motifs enriched in the promoters of (I) opening peaks with aging and (J) closing peaks with *Gdf3* deletion. See also Figure S4.

To predict potential TFs bound to accessible chromatin, we utilized footprinting analysis. Analyses of all identified ATAC-peaks (note that these were not differentially accessible peaks) revealed CTCF, PU.1, and JUN as the most prevalent TFs across all groups, irrespective of age or genotype (Figure S4H). This suggests that the key TFs in ATMs remain consistent regardless of these variables. To delve further, we predicted TFs on differentially accessible peaks using ShinyGO, a publicly available graphical gene-set enrichment tool^53^. Predictive analysis highlighted SMAD4 as the TF associated with opening peaks upon aging, and similarly, with closing peaks upon *Gdf3* deletion in old ATMs, with the highest enrichment scores (Figure 3I and 3J). Importantly, SMAD4 is required for SMAD2/3 to translocate into nucleus and initiate transcription^54^. In summary, we demonstrate that *Gdf3* is necessary for age-related changes in chromatin accessibility and that ATMs’ phenotype may be mediated through the GDF3-SMAD2/3/4 signaling axis.

### Inhibition of SMAD3 results in reduced inflammatory macrophages and hypothermia during endotoxemia

We next hypothesized that SMAD2/3 is required for the inflammatory responses in an age-dependent manner. To test this hypothesis, young (4-months-old) and old (23-months-old) WT mice were injected with specific inhibitor for SMAD3 (SIS3) and challenged with 0.1mg/kg LPS (Figure 4A)^55^. First, we validated the efficacy of SIS3 by quantifying pSMAD2/3 via phospho-flow. SIS3 effectively dampened pSMAD2/3 MFI and pSMAD2/3^+^ myeloid cells and ATMs from old mice (Figure 4B-D). However, SIS3 had no effect on lymphocytes (gated based on size), which exhibited lower frequency of pSMAD2/3^+^ cells than myeloid cells at a basal level (Figure 4B). Moreover, canonical SMAD2/3-transcribed genes (*Fn1, Serpine1*) were downregulated in VAT from SIS3-injected young mice compared to vehicle-injected counterparts. Interestingly, this effect was absent in old mice, suggesting that SMAD2/3 may regulate a different set of genes in older individuals (Figure 4E and 4F).

**Figure 4.**
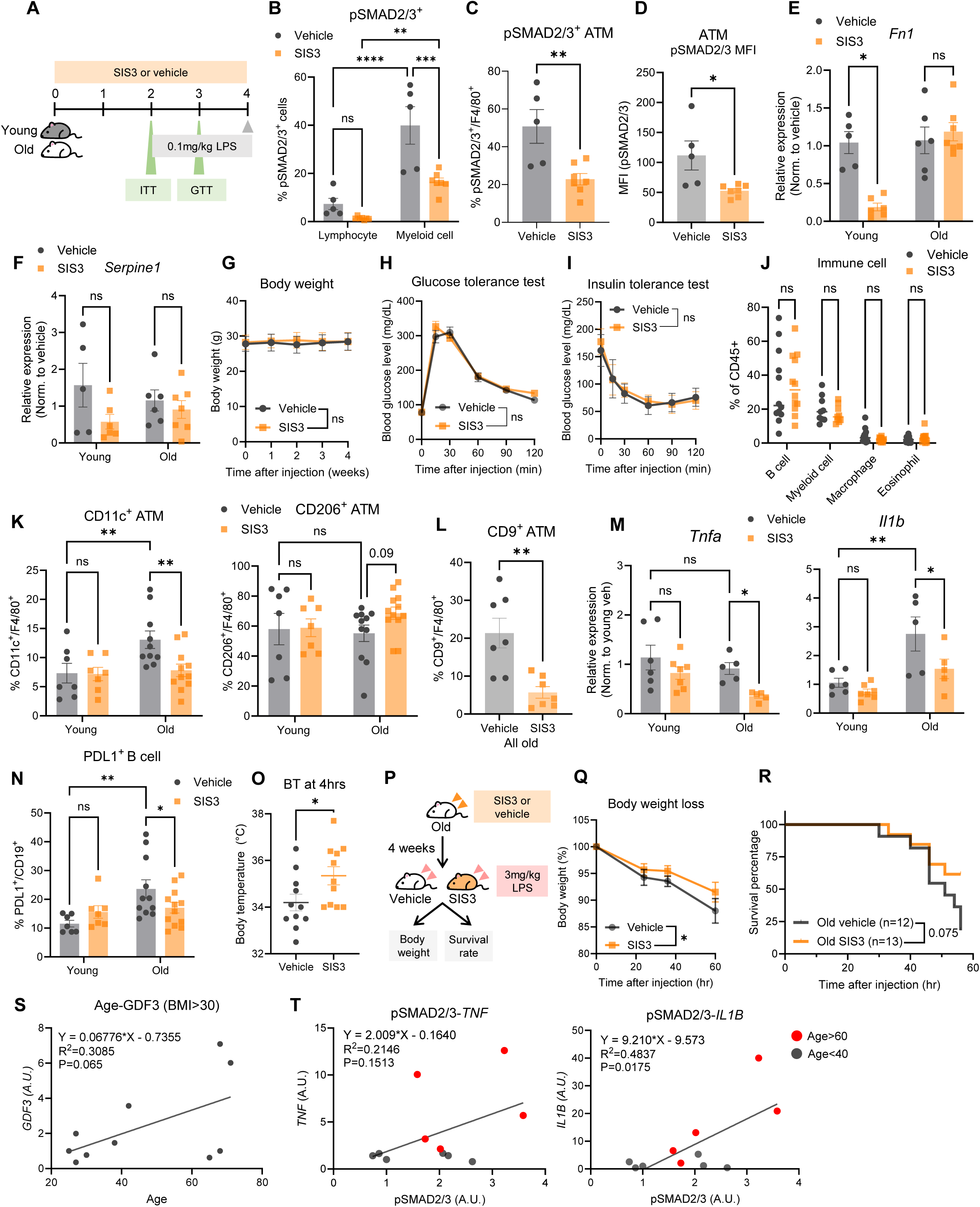
**Inhibition of SMAD3 alleviates inflammatory responses in old, but not young mice.** (A-O) Young (4-month-old) and old (23-month-old) female WT mice were i.p. injected with SIS3 or vehicle and LPS (0.1mg/kg) 4 hours prior to the end point. Young vehicle (YV), n=7; young SIS3 (YS), n=7; old vehicle (OV), n=11; old SIS3 (OS), n=11. (A) Schematic of experimental design. (B) Frequency of pSMAD2/3^+^ cells and (C) pSMAD2/3^+^ ATMs in VAT from old mice. OV, n=5; OS, n=7. (D) pSMAD2/3 MFI of ATMs from old mice. OV, n=5; OS, n=7. (E-F) VAT gene expression involved in fibrosis. YV, n=5; YS, n=6; OV, n=6; OS, n=7. (G) Body weights, (H) glucose sensitivity, and (I) insulin sensitivity of vehicle- and SIS3-injected old mice. (J) Frequency of immune cells, (K) CD11c^+^, CD206^+^ and (L) CD9^+^ ATMs in old VAT,. (M) VAT gene expression involved in inflammation. YV, n=6; YS, n=6; OV, n=5; OS, n=5. (N) Frequency of PDL1^+^ B cells in VAT. YV, n=7; YS, n=7; OV, n=11; OS, n=11. (O) Mean core BT of old mice post-LPS injection. OV, n=11; OS, n=11. (P-R) Old (20-month-old) female WT mice were injected with SIS3 or vehicle as shown above and LPS (3mg/kg). (P) Schematic of experimental design. (Q) Body weight loss (normalized to 0hr timepoint) after LPS injection. (R) Survival rate of vehicle or SIS3-injected mice after LPS injection. (S) Correlation between human VAT *GDF3* level and age. (T) Correlation between human VAT pSMAD2/3 level and inflammatory cytokine gene expression. All data are presented as means ± SEM. *p<0.05, **p<0.01, ***p<0.001, ****p<0.0001. All *in vivo* experiments were repeated independently twice. ns, not significant.

Acute blockade of SMAD3 signaling was shown to protect mice from diabetes and obesity with improved insulin or glucose sensitivity^31^. We asked if SIS3 would restore metabolic dysfunction that occurs during aging. However, month-long blockade of SMAD3 had no effect on the metabolic profile of old mice, shown by no differences in body weights, glucose sensitivity, and insulin sensitivity (Figure 4G-I). We next investigated whether inhibition of SMAD3 would reduce inflammatory responses in old mice, akin to phenotypes observed in *Gdf3*-deleted or JQ1-injected old mice. Overall immune composition showed no differences (Figure 4J). SIS3 reduced CD11c^+^ and CD9^+^ ATMs and increased CD206^+^ ATMs in old mice, but this effect was diminished in young mice (Figure 4K and 4L). Additionally, SMAD3 inhibition reduced inflammatory cytokine gene expression (*Tnfa, Il1b*) in old mice, but not in young mice (Figure 4M). Consistent with the effects of *Gdf3*-deletion on B cells, PDL1^+^ B cells were reduced in SIS3-injected old, but not young mice (Figure 4N). Inhibition of SMAD3 protected old mice from significant core BT loss observed in old mice injected with vehicle at 4 hours post-LPS (Figure 4O). Taken together, the data suggest that SMAD2/3 promotes inflammatory responses in an age-specific manner during endotoxemia.

Next, we sought to evaluate whether SMAD3 blockade, by decreasing inflammation, would alter the mortality upon endotoxemia induced by a lethal dose of LPS (3mg/kg, Figure 4P). We first monitored body weight loss as a non-invasive indicator of disease severity. As previously shown, body weights decreased with the severity of endotoxemia^56^. Notably, SIS3-injected old mice were partially protected from body weight loss compared to vehicle-injected mice (Figure 4Q). Moreover, SIS3-injected mice showed improved survival, although not statistically significant, in response to lethal endotoxemia (Figure 4R).

Reduced SMAD3 expression has been implicated in the longevity of humans^57^. Consistent with our observations in mouse models, *GDF3* expression was positively correlated with age in human VAT biopsies from bariatric surgeries (BMI>30; Figure 4S). Moreover, pSMAD2/3 positively correlated with inflammatory cytokine gene expression (*TNF, IL1B*), particularly in individuals aged over 60 years, further corroborating their role in inflammation in the aged (Figure 4T). Collectively, these data underscore the potential of SMAD2/3 inhibition for mitigating age-associated inflammation.

## DISCUSSION

Older individuals have increased risk for infections and subsequent sepsis, in part due to accumulating adiposity and a dysfunctional immune system. Gerotherapeutics that successfully improve the immune response are largely understudied. Our study reveals that the GDF3-SMAD2/3 axis may be a relevant pharmaceutical target. GDF3 promotes the inflammatory phenotype of ATMs in aged VAT, contributing to the exacerbation of endotoxemia-induced inflammation in older, but not younger, organisms. GDF3 signals through the SMAD2/3 pathway in ATMs and elicits pro-inflammatory responses diverging from SMAD2/3’s canonical immunoregulatory function^58,59^. Specifically, chromatin landscape of ATMs shifts towards inflammation upon aging, increasing the accessibility of inflammation-associated genes to the SMAD2/3/4 complex. ATAC-seq demonstrates that *Gdf3* deficiency can reverse the age-dependent changes in chromatin accessibility that is associated with the inflammatory phenotype of ATMs. Furthermore, genetic and pharmacological inhibition targeting the GDF3-SMAD2/3 axis reduces the inflammatory ATMs and protects against endotoxemia-induced inflammation and lethality in old mice.

The importance of VAT in aging and inflammation is corroborated by studies that demonstrate a beneficial effect of surgical removal of VAT on lifespan extension in a mouse model, as well as by studies that highlight the immunological role of VAT during metabolic challenge or infection^5–7,9,11,42,60^. Our study provides additional evidence for the role of GDF3 and VAT in endotoxemia during aging. Previous work indicates that IgG-mediated macrophage expression of *Tgfb* promotes fibrosis and metabolic decline via SMAD2/3 in aged VAT, suggesting potential functional overlap among TGFβ-family members^41^. Our data builds on this model, providing additional evidence for the importance of B cell-macrophage crosstalk, as lifelong deficiency of B cells reduces inflammatory macrophages^6^ and reduces *Gdf3* expression. We also provide evidence for the GDF3-SMAD2/3 axis regulating the phenotype of B cells, as indicated by a reduction in PDL1^+^ B cells in the genetically- and pharmacologically modified mouse models. Whether ligand-receptor interactions or exosome/organelle secretion could mediate this crosstalk is unknown. Our findings indicate that the mechanism governing inflammatory VAT microenvironment, driven by dysfunctional B cells and inflammatory macrophages, may converge on SMAD2/3 signaling.

Previous work highlights the altered chromatin state with inflammation, aging or obesity^32,35,47,61,62^. Our study complements these studies by identifying a significant increase in open chromatin peaks associated with inflammation in ATMs with aging. Consistent with GDF3 induction of pSMAD2/3 and inflammatory ATMs, predictive TF analysis points towards SMAD4 as a critical co-TF that could support the pro-inflammatory phenotype of ATMs. For instance, peaks associated with inflammatory cytokine receptors, such as *Il1r1* and *Il18r1,* become more accessible with aging and can be transcribed by the SMAD4 complex. SMAD2/3/4 regulate cell differentiation of embryonic stem cells by remodeling the chromatin^63,64^; however, our results demonstrate the use of this pathway to fine-tune the phenotype of macrophages. Moreover, our data show that inhibiting BRD4, an epigenetic modifier, reduces *Gdf3* and endotoxemia-related inflammation in old mice. Prior studies have revealed that BRD4 contributes to atherosclerosis-induced macrophage senescence^65^, as well as bacteria-induced macrophage chromatin remodeling^66^. These findings collectively suggest that BRD4 may play multiple roles during aging, promoting inflammation through both GDF3-independent and GDF3- dependent mechanisms^67^

In VAT, *Gdf3* expression is not induced by LPS treatment (data not shown), but recent work suggests that flexibility in the processing and release of the mature protein could mediate a rapid downstream response^68^. GDF3 would be expected to induce an immediate and local response. Previous studies show that GDF3 supports a pro-resolving phenotype of cardiac macrophages in young mice^20^. In contrast, our data indicates that GDF3 maintains inflammatory ATMs in old mice during endotoxemia. This context-dependency of GDF3 based on age of the organism further emphasizes the importance of the need for Gerotherapeutics. We quantified the GDF3 and pSMAD2/3 in adipose tissue from humans, which supports the relevance of the GDF3-SMAD2/3 axis in human disease and supports concepts that BRD4, GDF3 or SMAD2/3 could be a potential therapeutic target in older individuals. Whether chromatin remodeling is a required component for the therapeutic effect is unknown. Blockade of SMAD2/3 signaling through ALK inhibitors, proposed for treating multiple diseases including cancer, blood, and kidney diseases, shows promise in this context^69^. Inhibition of ALK7, the adipocyte-specific receptor, leads to a reduction in CD11c^+^ macrophages and the expression of *Il1b*, *Nlrp3* and *Gdf3* in VAT^70^. Lastly, inhibitors that target SMAD2/3 signaling, are currently under clinical development in phase I/II/III trials as cancer treatments^71^. Given these precedents, similar strategies to target the GDF3-SMAD2/3 could be taken to alleviate inflammation in the aging population.

## METHODS

### Animal care

All mice were housed in specific pathogen-free facilities in cage racks that deliver HEPA-filtered air to each cage with free access to sterile water at the University of Minnesota. Sentinel mice in our animal rooms were negative for currently tested standard murine pathogens at various times while the studies were performed. Female C57Bl6/J (WT) were bred from our colony, purchased from The Jackson Laboratory, or received from the National Institute of Aging Rodent colony. Littermate WT, *Gdf3*-/-, *Gdf3* fl/fl and *LysM-Cre* mice were bred and aged in our facility. Young mice were defined as 3- 6 months old and old mice were defined as 18-26 months old in all experiments. Mice were fed with normal chow diet (2018 Teklad Global 18% Protein Rodent Diets) and housed under 12hr light/dark cycles. All experiments and animal use were conducted in compliance with the National Institute of Health Guide for the Care and Use of Laboratory Animals and were approved by the Institutional Animal Care and Use Committee at the University of Minnesota. Old mice with extreme frailty, tumor, severe dermatitis, or other age-related pathologies were excluded from the study.

### Generation of *Gdf3* fl/fl and *Gdf3* -/- mice

Generation of *Gdf3* fl/fl mice was accomplished via CRISPR/Cas-mediated genome editing^72,73^. Potential Cas9 target guide (protospacer) sequences in intron 1 and 3’ were screened using the online tool CRISPOR^74^ and candidates were selected. Templates for sgRNA synthesis were generated by PCR from a pX330 template, sgRNAs were transcribed *in vitro* and purified (Megashortscript, MegaClear; ThermoFisher). sgRNA/Cas9 RNPs were complexed and tested for activity by zygote electroporation, followed by incubation of embryos to blastocyst stage, and genotype scoring of indel creation at the target sites. High-performing sgRNAs were selected, and repair template oligonucleotides (ssODN) incorporating loxP sites targeted to the Cas9 cleavage sites were synthesized by IDT. Guide RNA (gRNA) sequences are as follows: intron 1, 5’ guide: TATGAGGGAGGCTACTTAGG and 3’ guide CTAGGCTAGGAGTGTGCTTA. Guide primers for generating the template for transcription included a 5’ T7 promoter and a 3’ sgRNA scaffold sequence and were as follows (protospacer sequence underlined): 5’*TGTAATACGACTCACTATAGG*<u>TATGAGGGAGGCTACTTAGG</u>*GTTTTAGAGCTAGAAATAGC* 3’ 5’*TGTAATACGACTCACTATAGG*<u>CTAGGCTAGGAGTGTGCTTA</u>*GTTTTAGAGCTAGAAATAGC* 3’ sgRNA template reverse primer: 5’ AAAAGCACCGACTCGGTGCC 3’ Recombination templates: *Gdf3* 5’loxP ssODN: reverse strand 5’gcggaggcagagctggaggtatgagggaggctacttctcgag<u>ATAACTTCGTATAGCATACATTATACGAAGTTAT</u>aggaggtgtgtgtgtgtgt gtgtgtgtgtgtgtgtgtgtgtgtgtgtgtgtgtgtatacgtgtacgtgtatgcgcgcgtgcgctcgtgggt 3’ *Gdf3* 3’loxP ssODN: reverse strand 5’ggtgtgggtagtctcgggactaggctaggagtgtgc<u>ATAACTTCGTATAGCATACATTATACGAAGTTAT</u>gtcgacttagggtaaatcctttaataa aactaccaccccacctttggcttatgctctcttgaatcggacatgtctgtcatcatggttcatgagcccc

The floxed allele was created in two steps: targeting the 3’ loxP site, followed by a generation of breeding and subsequently targeting the 5’ loxP site. sgRNA/Cas9 RNP and the corresponding template oligo were electroporated into C57Bl/6J zygotes (The Jackson Laboratory)^73^. Embryos were transferred to the oviducts of pseudopregnant CD-1 foster females using standard techniques. Genotype screening of tissue biopsies from founder pups was performed by PCR amplification and Sanger sequencing to identify the desired alleles, including PCR to span both loxP sites. This was followed by breeding and sequence confirmation to establish germline transmission of the correctly targeted allele. Homozygous *Gdf3*-/- mice were generated by first breeding *Gdf3 fl/fl (loxP/loxP)* mice with *β-actin-Cre* transgenic mice (B6.FVB-*Tmem163^Tg(ACTB-cre)2Mrt^*/EmsJ, The Jackson Laboratory) and a second cross back to *Gdf3* fl/fl mice. This resulted in germline deletion of *Gdf3* that was used to generate littermate *Gdf3*+/+ and *Gdf3*-/- littermate controls.

The primers used to identify genotype are: 3F2 (TCGCCCATCTCCATGCTCTA), 3R2 (TAAGCTCACCAAGGGGTCCA), 5F3 (GAAGAGGAAGCTGATGCTGAC), and 5R3 (CTTCAGGGGCACTAGGCAAT).

### Mouse models

**For glucose tolerance test (GTT)**: Mice were fasted overnight for 16hr and intraperitoneally (i.p.) injected with 1.5mg/kg glucose. Blood glucose levels were measured at 0-, 30-, 60-, 90- and 120-min timepoint using glucometer.

**For insulin tolerance test (ITT)**: Mice were fasted for 4hr and i.p. injected with 0.8U/kg insulin. Blood glucose levels were measured at 0-, 30-, 60-, 90- and 120-min timepoint using glucometer.

**For body composition measurement**: EchoMRI was used.

**For Lipopolysaccharide (LPS) injection**: Mice were i.p. injected with sterile phosphate buffered solution (PBS; Corning) or 0.1mg/kg LPS (*E. Coli* O111-B4; Sigma, L3024) diluted in PBS. Core BT was taken using a rectal thermometer at 0-, 2- and 4-hour timepoint after injection. Mice were euthanized by cervical dislocation under isoflurane anesthesia 4 hours after injection.

**For lethal dose LPS injection**: Mice were i.p. injected with 3mg/kg (young) or 18mg/kg (old) LPS diluted in PBS. Core BT was taken using a rectal thermometer at indicated timepoints.

**For *in vivo* cell proliferation study**: 40mg/ml stock solution of 5-ethynyl-2’-deoxyuridine (EdU; Santacruz, sc-284628A) was prepared by dissolving 50mg of EdU in 1.25ml of DMSO and stored in -20°C. Prior to injection, 200µl of EdU stock solution was diluted in 1.8ml of PBS. For vehicle, 200µl of DMSO was diluted in 1.8ml of PBS. Mice were i.p. injected with 25mg/kg of EdU daily for 4 consecutive days and euthanized on day 3.

### Bone marrow derived macrophages (BMDM) culture

Bone marrow cells were collected from femur and tibia. Once collected, cells were pelleted by centrifugation at 500xg for 5min and resuspended with 2ml of ACK lysing buffer to remove erythrocytes. After 2min of incubation at room temperature (RT), 5ml of RPMI supplemented with 10% FBS was added to quench the reaction. Then, cells were pelleted for counting purposes and seeded to 6-well plate (3million cells/well) or 12-well plate (1.5million cells/well) with RPMI supplemented with 10% FBS, 1% antibiotic-antimycotic and 25ng/mL M-CSF (R&D, LS004177). On day5, cells were supplemented with fresh 10ng/mL M-CSF. On day7, cells were washed with PBS and serum-starved overnight with RPMI supplemented with 1% antibiotic-antimycotic. On day8, cells were stimulated by 20ng/mL recombination GDF3 (rGDF3; R&D, 9009-GD) or indicated dose of JQ1 for 4 hours. Cells were then harvested for gene expression and flow cytometry analysis.

### Tissue digestion

Tissues were harvested and stored in RPMI with 10% FBS on ice prior to digestion. Visceral or subcutaneous adipose tissue was mechanically digested/minced with surgical scissors, then enzymatically digested in 0.1% collagenase II (Worthington, LS004174) in Hanks Buffered Salt Solution (Gibco, 14185052) for 30 min at 37°, vortexing every 10 min for 30min to 45min in the water bath set at 37°C with 800 rpm agitation and vortexed every 10min. Following digestion, cells were centrifuged at 1500rpm for 5min to obtain stromal vascular fraction (SVF, pellet) and adipocyte fraction (floating lipid layer). Adipocyte fraction was collected for gene expression. Erythrocytes were removed from SVF as described above. Then, SVF was washed and filtered through a 40µM filter for further use. Spleen was prepared using mechanical digestion only, erythrocytes were removed, washed, and filtered through 100µM and 40µM filter. Tissue from control and experimental groups were digested simultaneously.

### Staining for flow cytometry

For live/dead staining, cells were incubated with Ghost Dye Red 780 Viability Dye (Tonbo, 13-0865-T100) for 20min on ice, in the dark. Then, cells were washed and incubated with Fcblock (Thermo, 14-9161-73) and surface antibodies for 45min on ice, in the dark. For nuclear or intracellular antibodies, FoxP3/Transcription Factor Fix/Permeabilization kit (Invitrogen, 00-5521-00) and BD Cytofix/Cytoperm Fix/Permeabilization kit (BD, 554714) were used. For phospho-flow, cells were incubated with or Fixable Aqua Dead Cell Stain Kit (Invitrogen, L34957) for 10min on ice, in the dark. Then, cells were washed and incubated with warm BD Phosflow Fix Buffer (BD, 557870). After a 10min incubation at 37°C, cells were washed and permeabilized with ice-cold BD Phosflow Perm buffer III (BD, 558050) for 30min on ice, protected from light. Analysis was performed on FACSSymphony A3 cytometers using FlowJo v10; gating strategies are shown in supplemental figures. For fluorescence-activated cell sorting of ATM: Live CD45^+^ CD11b^+^ SiglecF^-^ F4/80^+^ cells from SVF were sorted on a BDFACSAriaII into RPMI with 20% FBS. Antibodies used are documented in Table S1.

### Single cell RNA sequencing analysis

#### Library preparation

Single-cell libraries were prepared in triplicate with >85% cell viability confirmed by trypan blue staining. Cells were resuspended in MEM with 10% FBS at 1200 cells/uL. Library construction used Chromium Next GEM Single Cell 3ʹ dual index Kits v3.1 with Chip G (10x Genomics). About 10,000 single cells were loaded to generate gel bead-in-emulsions (GEMs), lysed, and RNA was barcoded and reverse-transcribed. GEMs were broken, and cDNA was prepared following the manufacturer’s protocol. cDNA quality was assessed with the Agilent Bioanalyzer and Pico-green. Libraries underwent QC by shallow-sequencing on an Illumina NextSeq and final sequencing on an Illumina Nova Seq S4 capturing ∼20,000 paired-end reads per cell. Sequencing libraries were converted into feature-barcode matrices using Cell Ranger (10x Genomics, version 5.0.0) and mapped to GRCh38.

#### scRNA-seq QC and clustering

Analysis of read count matrices was performed using Seurat version 4.3.1 in R. Cells with mitochondrial RNA <15% and an R² of 0.95 for nCount_RNA vs. nFeature_RNA were included. Each sample was normalized using ‘NormalizeData’, and the top 2000 highly variable genes were selected with ‘FindVariableFeatures’. Dimensional reduction and clustering followed the standard Seurat workflow with ‘ScaleData’ and ‘RunPCA’. The top 50 principal components, identified by ‘ElbowPlot’, were used for UMAP visualization via ‘RunUMAP’ and finding nearest neighbors. Clusters and subclusters were identified using the Louvain algorithm with a resolution of 0.2.

#### scRNA-seq cluster identification and DEG list generation

Differentially expressed genes (DEGs) were identified by comparing normalized expressed RNAs for each model to RNAs expressed in control cells via the function ‘FindMarkers’ and then merging the results. When comparing clusters, the function ‘FindAllMarkers’ was employed to find marker genes that characterized a specific cluster compared to all other clusters. A minimum of 100 cells from a sample group in a cluster was required for downstream analysis.

### Western blotting

Visceral adipose tissue was snap frozen in liquid nitrogen immediately after harvest. Tissue was homogenized in RIPA buffer containing phosphatase inhibitor and protease inhibitors (Sigma-Aldrich, P0044, P5726, P8340). Lysates were left on ice for an hour and vortexed every 15min. Lysates were then centrifuged twice at 13,000rpm for 10min at 4°C. Protein concentration was quantified using the Bradford Protein Assay Kit (Thermo, 23246) and equal amounts of protein were run on an 4-12% Bis-Tris SDS-PAGE gel and transferred to PVDF membrane via semi-dry transfer method. Blots were probed with primary antibodies and then incubated in secondary antibodies of appropriate species. Then, Femto Maximum Sensitivity Substrate (Thermo, 34096) or ECL Western Blotting Substrate (Thermo, 32209) was used for detection. Quantification of bands was done with Thermofisher’s iBright analysis software.

### RNA extraction and gene expression analysis

<u>Whole tissues</u> were homogenized in Trizol (Invitrogen,15596026) using the Next Advantage Bullet Blender Storm 24. Following homogenization, chloroform was added to samples and incubated at RT for 15min. Samples were centrifuged at 13,000rpm for 15min at 4°C. Then, 70% molecular grade ethanol was added to aqueous phase of centrifuged homogenate for RNA extraction using PureLink RNA Mini Kits (Invitrogen, 12183025) according to manufacturer’s instructions. <u>For suspended cells</u>, they were lysed in lysis buffer supplemented with 1% β-mercaptoethanol, incubated and then mixed with 70% molecular grade ethanol. Reverse transcription PCR and quantitative PCR were performed as previously described. Primer sequences used for qPCR are shown in Table S2.

### Adipose tissue macrophage culture

ATMs were positively selected from freshly collected SVF using EasySep Mouse F4/80 Positive Selection Kit (Stemcell, 100-0659) according to manufacturer’s instructions. After selection, cells were incubated in RPMI with 10% FBS and 1% antibiotic-antimycotic and stimulated with vehicle or rGDF3 (20ng/mL) for 30 minutes for flow cytometry analysis.

### Ex vivo lipolysis

Visceral adipose tissues were collected and weighed, ranging from 15mg to 20mg. Then, tissues were transferred to 96-well plate and cultured in 100µl lipolysis buffer (Krebs buffer with 3.5% fatty acid free BSA and 0.1% glucose). The buffer was collected after a 2hr incubation at 37°C for Non-esterified Fatty Acid assay (Fujifilm HR Series), following the manufacturer’s instructions. The NEFA values calculated from standard curve were then normalized to tissue weights.

### Human tissues

Frozen visceral adipose tissues were obtained from bariatric surgeries performed at the University of Minnesota. The detailed age (sex, BMI) information is as follows: 25 (F, 39.26), 27 (F, 39.49), 27(F, 38.74), 30 (F, 39.84), 38 (F, 52.5), 42 (F, 37), 65 (F, 35.4), 66, (F, 18.8), 66, (F, 28.5), 68 (F, 41.1), 68 (F, 22), 71 (F, 37.2).

### Statistical analysis

Statistical significance was calculated using a multiple t-test, unpaired Student’s t-test, paired t-test or ANOVA with post-hoc tests for multiple comparisons. *P < 0.05; **P < 0.005; ***P < 0.001; ****P < 0.0001. One-way ANOVA was used when there was one experimental condition, but multiple groups (i.e. pSMAD2/3 MFI comparison between immune cells) and two-way ANOVA was used when there were two experimental conditions (i.e. age and genotype). Paired t-test was used to compare the effects of a treatment on the cells derived from the same individual or sample with different treatment conditions (i.e. vehicle or rGDF3 treatment on ATMs). For a survival experiment, the Log-rank (Mantel-Cox) test was used to determine statistical significance. For correlations, Pearson’s correlation analysis was used. Statistical outliers were identified and excluded using GraphPad Prism. A 95% confidence interval was used for all tests. Data were assumed to be normally distributed unless standard deviations significantly differed between groups. Sample sizes were based on previous experiments. All tests were performed using GraphPad Prism v10. Data are expressed as mean ± S.E.M. Details of biological replicates are in the figure legends.

### ATAC-seq library preparation

ATAC-seq kit (Activemotif, 53150) was used for ATAC-seq library preparation. In brief, ATMs (live CD45^+^ CD11b^+^ SiglecF^-^ F4/80^+^ cells) were sorted using a BDFACSAriaII instrument. Cells were washed with ice-cold PBS to remove any debris, and 10^6^ cells were resuspended in 100μl ice-cold ATAC lysis buffer. Then, cells were gently pipetted up and down and incubated on ice for 5min. These cells were centrifuged at 500xg for 10min at 4°C and supernatant was discarded. Next, 50μl of tagmentation master mix containing buffer, 10xPBS, 1% Digitonin, 10% Tween 20, H2O and assembled transposome was added to each sample. Samples were incubated at 37°C for 30 min in a thermomixer set at 800rpm. Then, tagmented DNA was extracted and amplified (72°C for 5min; 98°C for 30 sec; 12 cycles of 98°C for 10sec, 63°C for 30sec, 72°C for 1min) for library preparation. Lastly, 1.2x SPRI bead solution was added to samples for left side size selection. The samples were frozen and sent to Novogene (CA), sequenced and analyzed.

### ATAC bioinformatics analysis

First, FastQC was used to assess the quality of the raw reads. Then, Skewer^75^ was used to trim and filter off the sequencing adaptors and low-quality bases (mean quality>20, N ratio<15%, post-trimming length>18nt), and trimmed reads were mapped to mouse genome GRCm39 (mm39) using BWA ^76^. After alignment, MACS2 software^77^ was used with 100 bp shift, 200 bp extension for peak calling analysis. Only peaks called with a peak score (or q-value) of 5% or better were kept from each sample and the number of peaks, the peak width, its distribution, and the peak related genes were determined with MACS2. Moreover, Homer software was used for motif enrichment analysis. Consensus sequences were identified by using the sequence of 250pb (total 500bp) upstream and downstream of the peak. To predict the accessibility around transcription start site (TSS), PeakAnalyzer^78^ was utilized to analyze Peak-TSS distance distribution. The distribution of peaks in functional area was analyzed by ChIPseeker^79^. Differential peaks identified and visualized by edgeR^80^ (Fold Change>2, FDR<0.05) were analyzed using ShinyGo 0.80^53^ for transcription factor predictions and gene-set enrichment analyses, including KEGG and GO. For visualization of mapped reads, Integrative Genomics Viewer analyzer^81^ was used.

## Acknowledgements

We acknowledge and thank the University of Minnesota Flow Cytometry Resource, specifically Paul Champoux and Rashi Arora, for their expertise. We thank the Yale Genome Editing Center staff for their help in generating the *Gdf3* cKO mouse and UMN Research Animal Resources Staff for their day-to-day care of mice.

## Author Contributions

Conceptualization: IJ, CDC; Investigation: IJ, VK, SC, DS, AC, DAB, CDC; Data curation – formal analysis: IJ, CDC; scRNA sequencing data analysis: AC; Mouse model generation: SB, TN; Project administration: IJ, CDC; Oversight: CDC; Writing – original draft: IJ, CDC; Writing – review, editing, and revision: All authors; Funding acquisition: CDC, DAB

## Material & Correspondence

Correspondence and material requests should be addressed to C.D.C.

## Data availability

Raw and processed single-cell RNA sequencing data are available at Gene Expression Omnibus (GEO) GSE274935. ATAC sequencing data are available at the GEO <u>GSE272899</u> – to be released when published.

---

**Figure S1.**
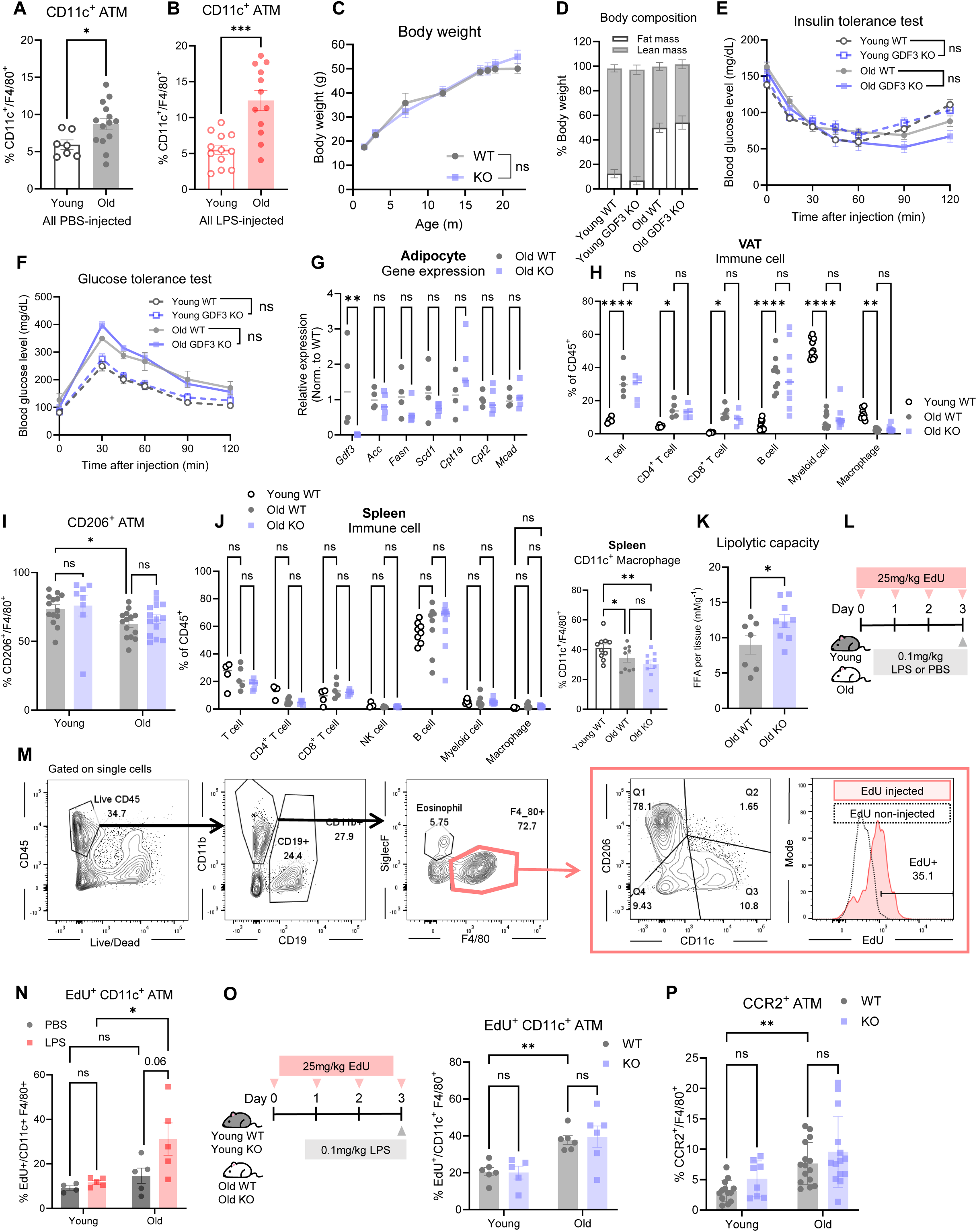
Lifelong *Gdf3* deficiency does not affect metabolic profile of old mice, nor the proliferation or infiltration of ATMs in old VAT, related to Figure 1. (A-B) Frequency of CD11c^+^ ATMs in VAT from young (n=7) or old (n=15) female WT mice injected with (A) PBS or (B) 0.1mg/kg LPS. (C) Body weights of female WT (n=11) and GDF3 KO (n=8) mice measured throughout lifetime. (D) Body composition of YW (n=6), YK (n=3), OW (n=8), and OK (n=8) mice. (E) Insulin tolerance test and (F) glucose tolerance test of 2∼3- and 18∼19-month WT and KO mice. YW, n=5; YK, n=5; OW, n=14; OK, n=11. (G-K) Young (3- to 4-month-old) WT, old (19- to 22-month-old) WT or KO mice were challenged with 0.1mg/kg LPS. (G) Adipocyte gene expression involved in fatty acid oxidation and fatty acid synthesis. OW, n=4; OK, n=6. (H) Frequency of immune cells (gated as % of CD45^+^) in VAT. YW, n=4∼10; OW, n=5∼10; OK, n=5∼8. (I) Frequency of CD206^+^ ATMs. YW, n=14; YK, n=8; OW, n=15; OK, n=14. (J) Frequency of immune cells (gated as % of CD45^+^) in spleen. YW, n=4∼10; OW, n=5∼10; OK, n=5∼8. (K) Free fatty acid released from VAT explants. OW, n=7; OK, n=9. (L-O) Young (4-months-old) and old (24-months-old) female WT mice we injected with 25mg/kg EdU for 4 consecutive days and 0.1mg/kg LPS or PBS for 4 hours on day 3. (L) Experimental schematic of *in vivo* proliferation assay. (M) Representative gating strategy. (N) Frequency of EdU^+^CD11c^+^ ATMs in VAT. n=5 per group. (O) Schematic of experimental design (left). Frequency of EdU^+^CD11c^+^ ATMs in VAT (right). YW, n=6; YK, n=5; OW, n=6; OK, n=6. (P) Frequency of CCR2^+^ ATMs in VAT. YW, n=14; YK, n=8; OW, n=15; OK, n=14 All data are presented as means ± SEM.

**Figure S2.**
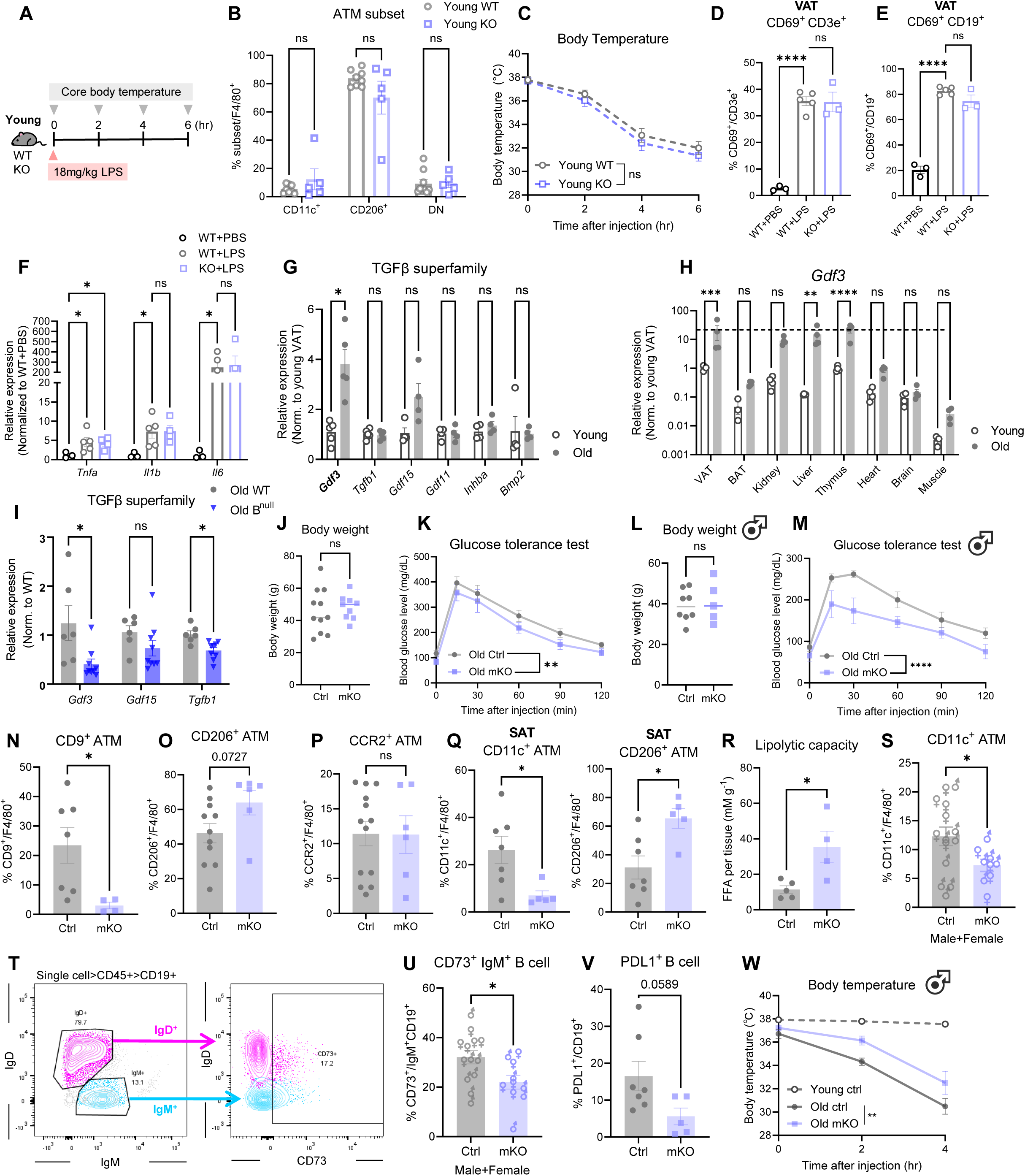
*Gdf3* is not required for LPS-induced lethality in young mice but is regulated with age in a B cell-dependent manner, related to Figure 1. (A-F) Young (4- to 6-months-old) female WT and KO mice were injected with lethal dose of LPS (18mg/kg) and VAT was collected for analysis 6hr post-LPS injection. (A) Schematic of experimental design. (B) Frequency of ATMs’ subpopulation in VAT. WT, n=8; KO, n=5. (C) Mean core BT after LPS injection. WT, n=13; KO, n=8. (D-E) Frequency of (D) activated T cells and (E) activated B cells in VAT. WT+PBS, n=3; WT+LPS, n=5; KO+LPS, n=4. (F) Inflammatory gene expression in VAT. WT+PBS, n=3; WT+LPS, n=5; KO+LPS, n=4. (G) TGFβ superfamily gene expression in VAT and (H) *Gdf3* expression in multiple tissues and from 2-months-old (n=4) and 21-month-old (n=4) female WT mice. (I) TGFβ superfamily gene expression in VAT from 20- to 22-month-old female WT (n=6) and B^null^ (n=9) mice injected with 0.1mg/kg LPS. (J-M) Metabolic profiling of old (21- to 26-month-old) control and mKO mice. Female control, n=11; female mKO, n=9; male control, n=8; male mKO, n=5. Body weights and glucose tolerance test of female (J-K) and male (L-M) mice. (N-R) Old (21- to 24-month-old) female control and mKO mice were injected with 0.1mg/kg LPS. Frequency of (N) CD9^+^ ATMs, (O) CD206^+^ ATMs, and (P) CCR2^+^ ATMs in VAT. (Q) Frequency of CD11c^+^ and CD206^+^ ATMs in SAT. Control, n=7∼14; mKO, n=4∼6. (R) Free fatty acid released from VAT explants from old female control (n=5) and mKO (n=4) mice. (S) Frequency of CD11c^+^ ATMs from VAT of male and female mice injected with 0.1mg/kg LPS. Females ♀, males ♂. Control, n=14; mKO, n=9. (T) Representative gating strategy for B cells. (U-V) Frequency of (U) CD73^+^IgM^+^ B cells and (V) PDL1^+^ B cells in VAT of old control and mKO mice injected with 0.1mg/kg LPS. (W) Mean core BT of young (3-month-old) or old (23- to 26-month-old) male mice post-LPS (0.1mg/kg) injection. Young control, n=8; old control n=8; old mKO, n=5. All data are presented as means ± SEM.

**Figure S3.**
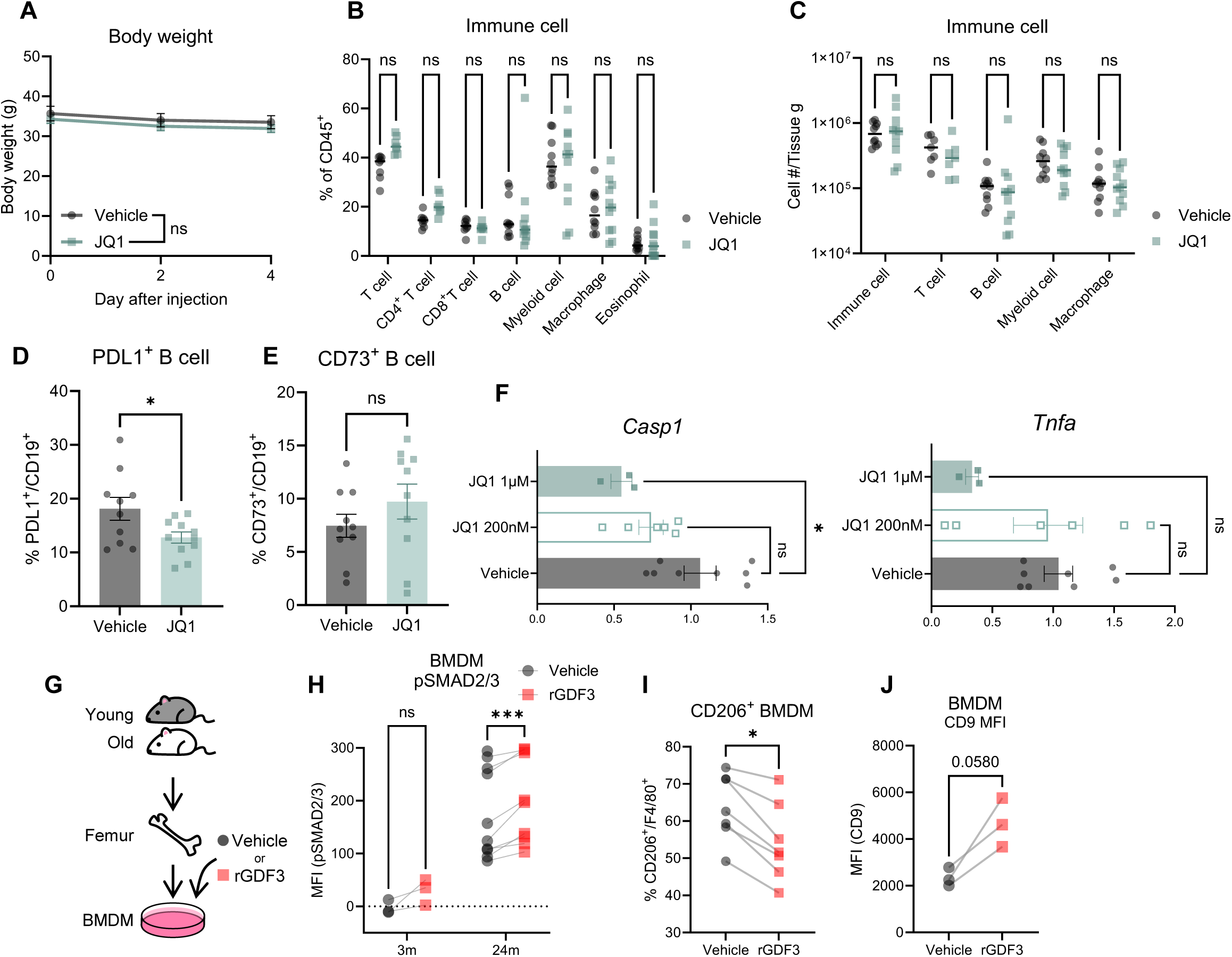
BRD4-regulated *Gdf3* promotes pSMAD2/3, related to Figure 2. (A) Body weights of vehicle- or JQ1-injected old (23- to 24-month) female mice. Vehicle-injected, n=10; JQ1-injected, n=11. (B-H) Old (23- to 24-month-old) female WT mice were i.p. injected with JQ1 or vehicle and LPS (0.1mg/kg). Vehicle-injected, n=10; JQ1-injected, n=11. (B) Frequency and (C) cellularity of immune cells in VAT from vehicle- or JQ1-injected old mice. (D-E) Frequency of (D) PDL1^+^ and (E) CD73^+^ B cells in VAT. (F) *Casp1* and *Tnfa* expression in BMDMs treated with vehicle or JQ1. (G-H) BMDMs from 3- or 24-month WT mice were treated with vehicle or rGDF3 (20ng/mL) *in vitro*. (G) Schematic of experimental design for BMDMs. (H) pSMAD2/3 MFI in BMDMs. (I) Frequency of CD206^+^ BMDMs and (J) MFI of CD9 of BMDMs. All data are presented as means ± SEM.

**Figure S4.**
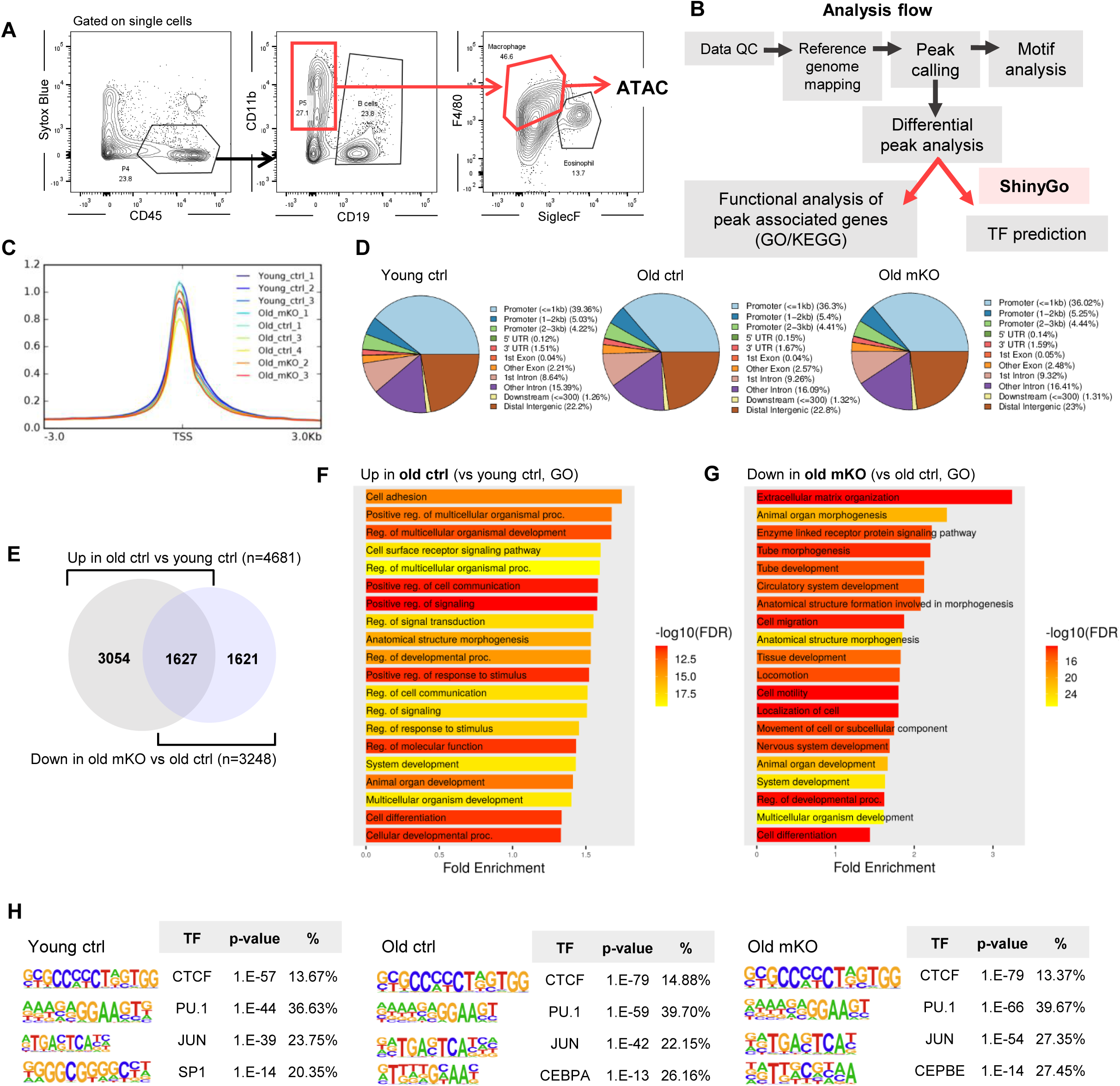
ATAC sequencing of ATMs, related to. Figure 3. (A) Representative gating strategy for fluorescence-activated cell sorting (FACS). (B) Schematic of bioinformatics analysis flow. (C) Transcription start site (TSS) enrichment plot showing the distribution of the reads mapped to TSS. (D) Representative peak annotation pie charts for identified ATAC-seq peaks. (E) Ven diagram depicting the overlap between opening peaks with aging (old vs young control) and closing peaks with *Gdf3* deletion (old mKO vs old control) from ATAC-seq analysis. (F-G) Gene ontology (GO Biological Process) analysis of genes associated with (F) opening peaks with aging and (G) closing peaks with *Gdf3* deletion. (H) De novo motif search within all identified ATAC-seq peaks from young ctrl, old ctrl and old mKO ATMs. Consensus motif, transcription factor, p-value, and percentage of targets are shown in table. * ATAC-seq was performed in 3 biological replicates (young control, n=3; old control, n=3; old mKO, n=3).

**Supplementary Table 1.**
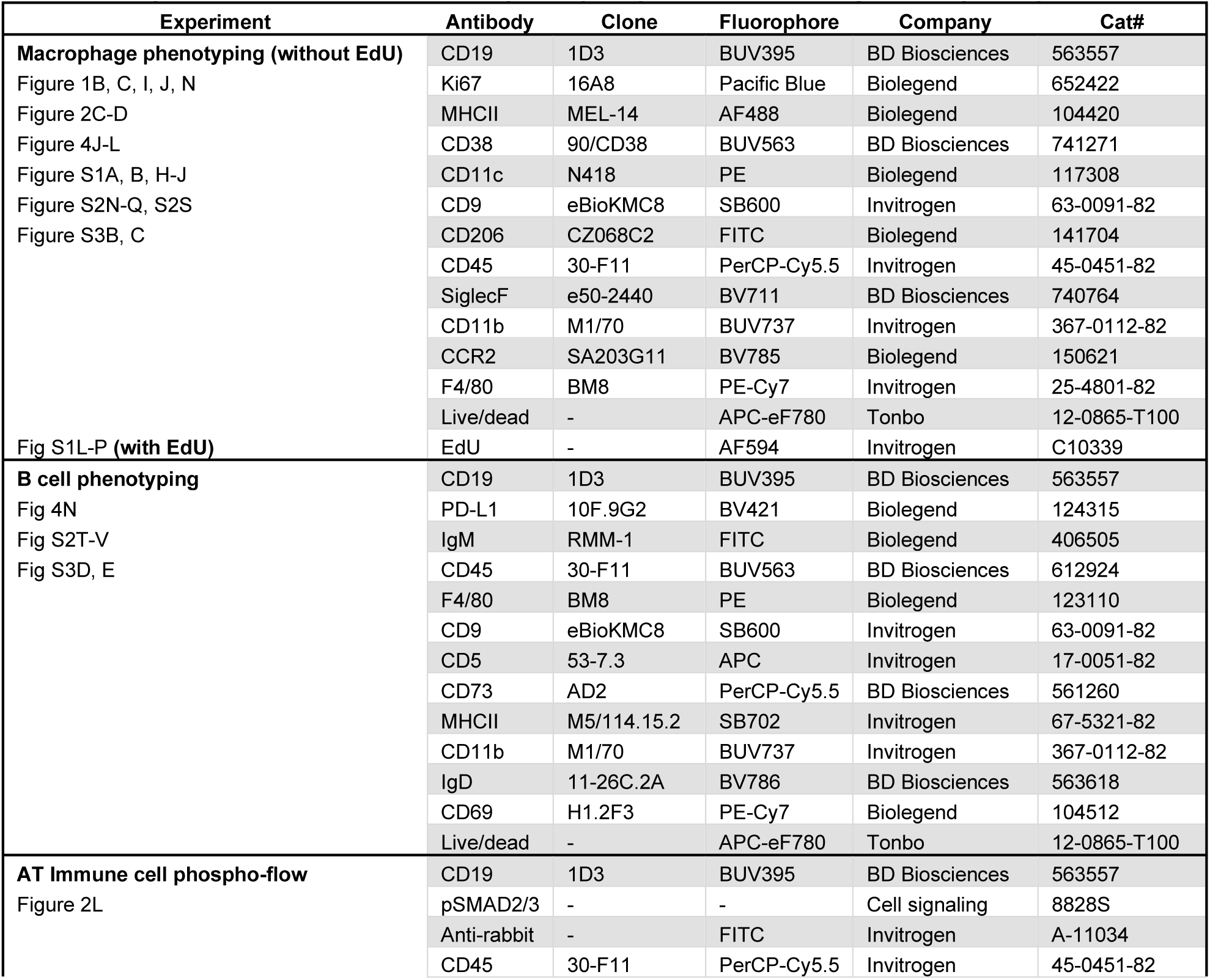
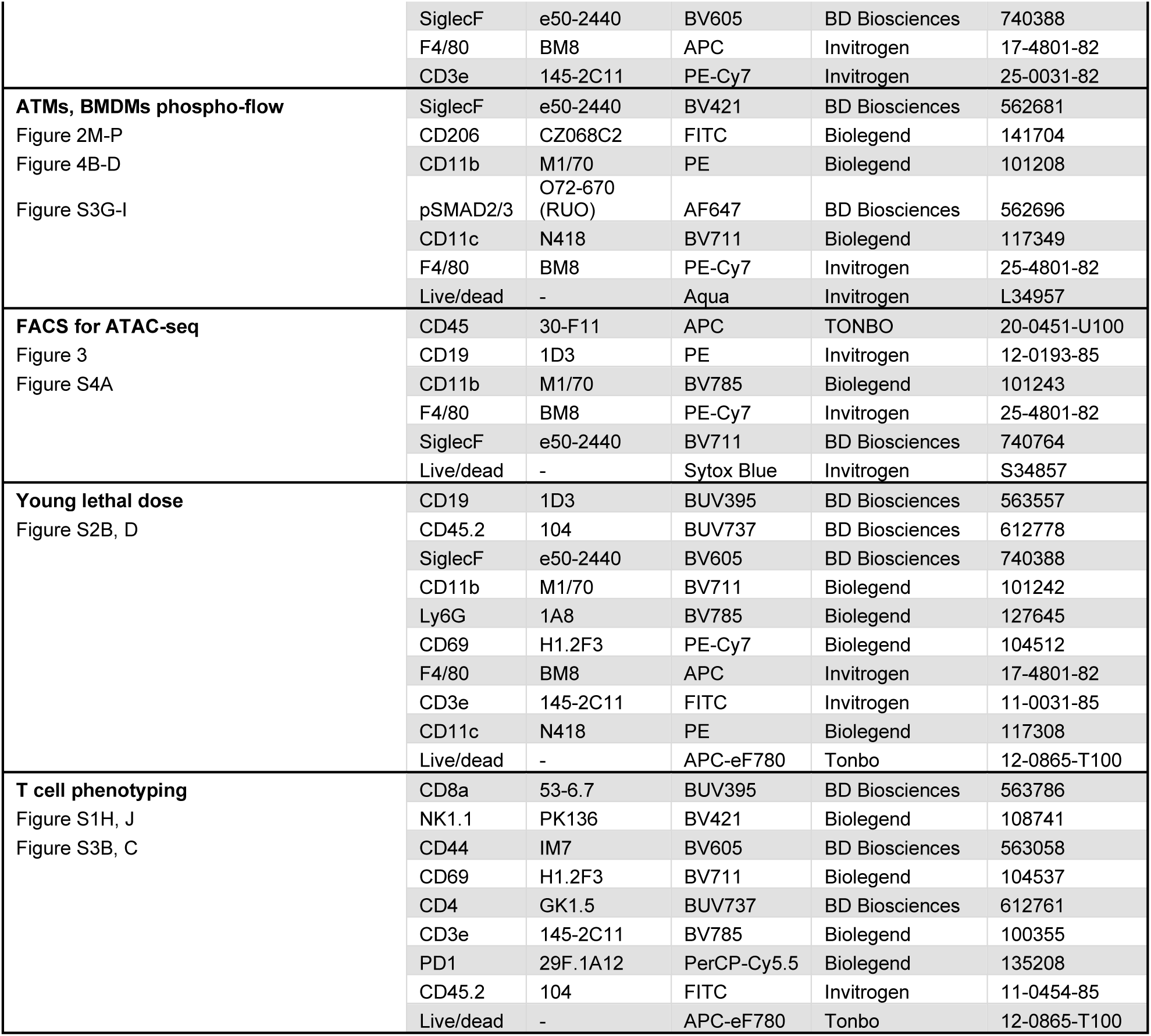
Antibodies used for flow cytometry analysis, related to staining for flow cytometry

| Experiment | Antibody | Clone | Fluorophore | Company | Cat# |
| --- | --- | --- | --- | --- | --- |
| **Macrophage phenotyping (without EdU)**<br>Figure 1B, C, I, J, N<br>Figure 2C-D<br>Figure 4J-L<br>Figure S1A, B, H-J<br>Figure S2N-Q, S2S<br>Figure S3B, C | CD19 | 1D3 | BUV395 | BD Biosciences | 563557 |
|  | Ki67 | 16A8 | Pacific Blue | Biolegend | 652422 |
|  | MHCII | MEL-14 | AF488 | Biolegend | 104420 |
|  | CD38 | 90/CD38 | BUV563 | BD Biosciences | 741271 |
|  | CD11c | N418 | PE | Biolegend | 117308 |
|  | CD9 | eBioKMC8 | SB600 | Invitrogen | 63-0091-82 |
|  | CD206 | CZ068C2 | FITC | Biolegend | 141704 |
|  | CD45 | 30-F11 | PerCP-Cy5.5 | Invitrogen | 45-0451-82 |
|  | SiglecF | e50-2440 | BV711 | BD Biosciences | 740764 |
|  | CD11b | M1/70 | BUV737 | Invitrogen | 367-0112-82 |
|  | CCR2 | SA203G11 | BV785 | Biolegend | 150621 |
|  | F4/80 | BM8 | PE-Cy7 | Invitrogen | 25-4801-82 |
|  | Live/dead | - | APC-eF780 | Tonbo | 12-0865-T100 |
| **Fig S1L-P (with EdU)** | EdU | - | AF594 | Invitrogen | C10339 |
| **B cell phenotyping**<br>Fig 4N<br>Fig S2T-V<br>Fig S3D, E | CD19 | 1D3 | BUV395 | BD Biosciences | 563557 |
|  | PD-L1 | 10F.9G2 | BV421 | Biolegend | 124315 |
|  | IgM | RMM-1 | FITC | Biolegend | 406505 |
|  | CD45 | 30-F11 | BUV563 | BD Biosciences | 612924 |
|  | F4/80 | BM8 | PE | Biolegend | 123110 |
|  | CD9 | eBioKMC8 | SB600 | Invitrogen | 63-0091-82 |
|  | CD5 | 53-7.3 | APC | Invitrogen | 17-0051-82 |
|  | CD73 | AD2 | PerCP-Cy5.5 | BD Biosciences | 561260 |
|  | MHCII | M5/114.15.2 | SB702 | Invitrogen | 67-5321-82 |
|  | CD11b | M1/70 | BUV737 | Invitrogen | 367-0112-82 |
|  | IgD | 11-26C.2A | BV786 | BD Biosciences | 563618 |
|  | CD69 | H1.2F3 | PE-Cy7 | Biolegend | 104512 |
|  | Live/dead | - | APC-eF780 | Tonbo | 12-0865-T100 |
| **AT Immune cell phospho-flow**<br>Figure 2L | CD19 | 1D3 | BUV395 | BD Biosciences | 563557 |
|  | pSMAD2/3 | - | - | Cell signaling | 8828S |
|  | Anti-rabbit | - | FITC | Invitrogen | A-11034 |
|  | CD45 | 30-F11 | PerCP-Cy5.5 | Invitrogen | 45-0451-82 |
|  | SiglecF | e50-2440 | BV605 | BD Biosciences | 740388 |
|  | F4/80 | BM8 | APC | Invitrogen | 17-4801-82 |
|  | CD3e | 145-2C11 | PE-Cy7 | Invitrogen | 25-0031-82 |
| **ATMs, BMDMs phospho-flow**<br>Figure 2M-P<br>Figure 4B-D<br>Figure S3G-I | SiglecF | e50-2440 | BV421 | BD Biosciences | 562681 |
|  | CD206 | CZ068C2 | FITC | Biolegend | 141704 |
|  | CD11b | M1/70 | PE | Biolegend | 101208 |
|  | pSMAD2/3 | O72-670 (RUO) | AF647 | BD Biosciences | 562696 |
|  | CD11c | N418 | BV711 | Biolegend | 117349 |
|  | F4/80 | BM8 | PE-Cy7 | Invitrogen | 25-4801-82 |
|  | Live/dead | - | Aqua | Invitrogen | L34957 |
| **FACS for ATAC-seq**<br>Figure 3<br>Figure S4A | CD45 | 30-F11 | APC | TONBO | 20-0451-U100 |
|  | CD19 | 1D3 | PE | Invitrogen | 12-0193-85 |
|  | CD11b | M1/70 | BV785 | Biolegend | 101243 |
|  | F4/80 | BM8 | PE-Cy7 | Invitrogen | 25-4801-82 |
|  | SiglecF | e50-2440 | BV711 | BD Biosciences | 740764 |
|  | Live/dead | - | Sytox Blue | Invitrogen | S34857 |
| **Young lethal dose**<br>Figure S2B, D | CD19 | 1D3 | BUV395 | BD Biosciences | 563557 |
|  | CD45.2 | 104 | BUV737 | BD Biosciences | 612778 |
|  | SiglecF | e50-2440 | BV605 | BD Biosciences | 740388 |
|  | CD11b | M1/70 | BV711 | Biolegend | 101242 |
|  | Ly6G | 1A8 | BV785 | Biolegend | 127645 |
|  | CD69 | H1.2F3 | PE-Cy7 | Biolegend | 104512 |
|  | F4/80 | BM8 | APC | Invitrogen | 17-4801-82 |
|  | CD3e | 145-2C11 | FITC | Invitrogen | 11-0031-85 |
|  | CD11c | N418 | PE | Biolegend | 117308 |
|  | Live/dead | - | APC-eF780 | Tonbo | 12-0865-T100 |
| **T cell phenotyping**<br>Figure S1H, J<br>Figure S3B, C | CD8a | 53-6.7 | BUV395 | BD Biosciences | 563786 |
|  | NK1.1 | PK136 | BV421 | Biolegend | 108741 |
|  | CD44 | IM7 | BV605 | BD Biosciences | 563058 |
|  | CD69 | H1.2F3 | BV711 | Biolegend | 104537 |
|  | CD4 | GK1.5 | BUV737 | BD Biosciences | 612761 |
|  | CD3e | 145-2C11 | BV785 | Biolegend | 100355 |
|  | PD1 | 29F.1A12 | PerCP-Cy5.5 | Biolegend | 135208 |
|  | CD45.2 | 104 | FITC | Invitrogen | 11-0454-85 |
|  | Live/dead | - | APC-eF780 | Tonbo | 12-0865-T100 |

**Supplementary Table 2.**
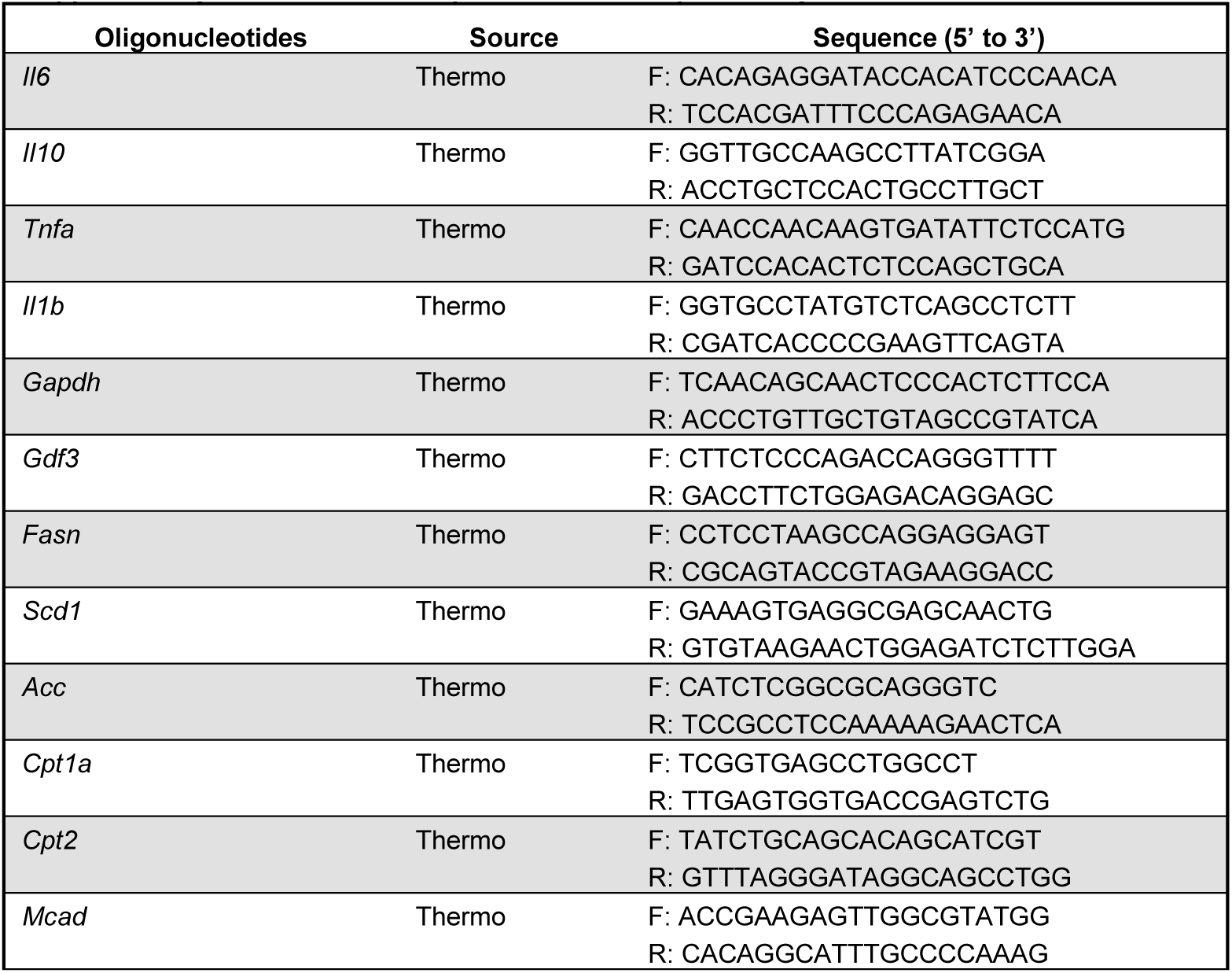
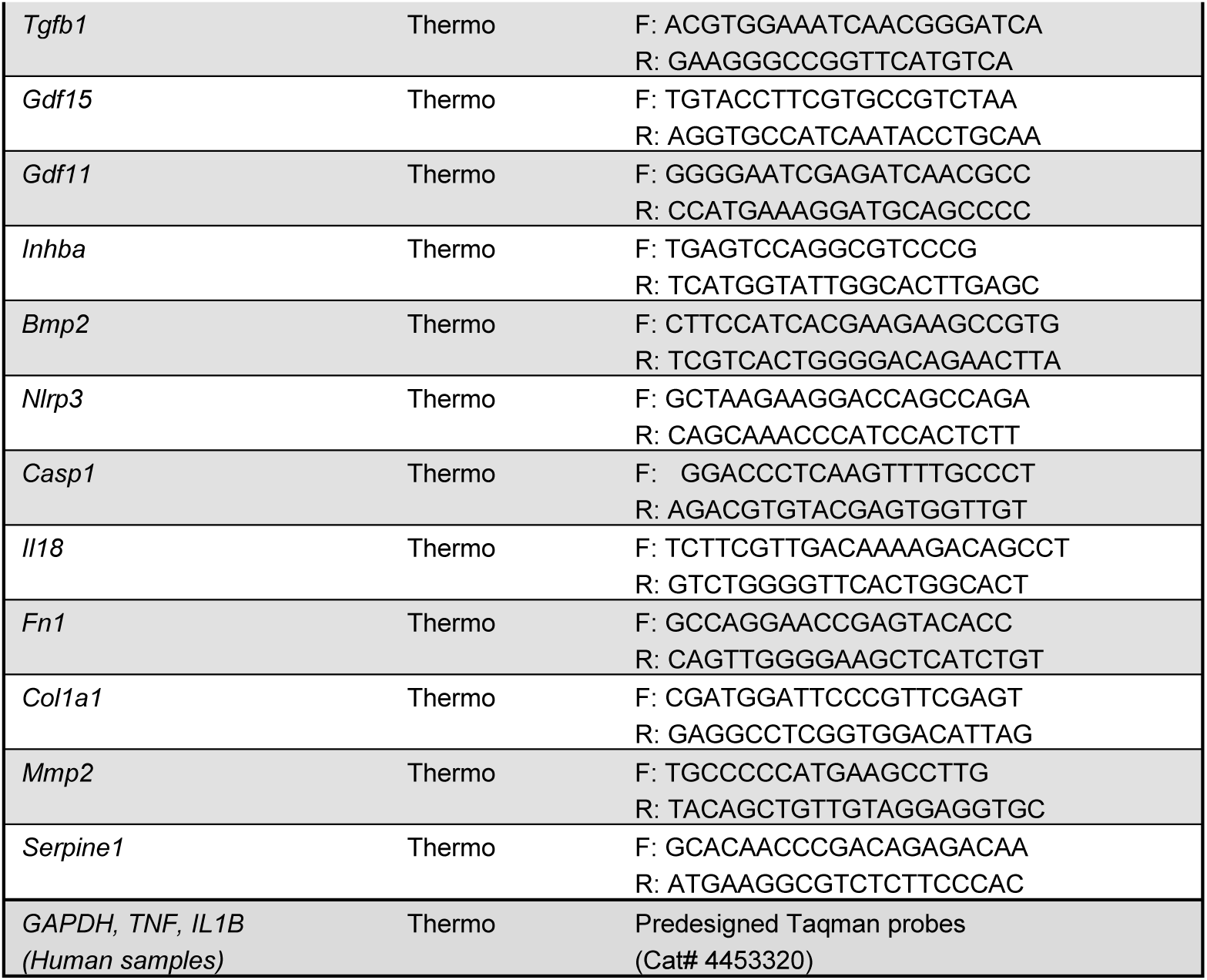
Primer sequences used for qPCR analysis

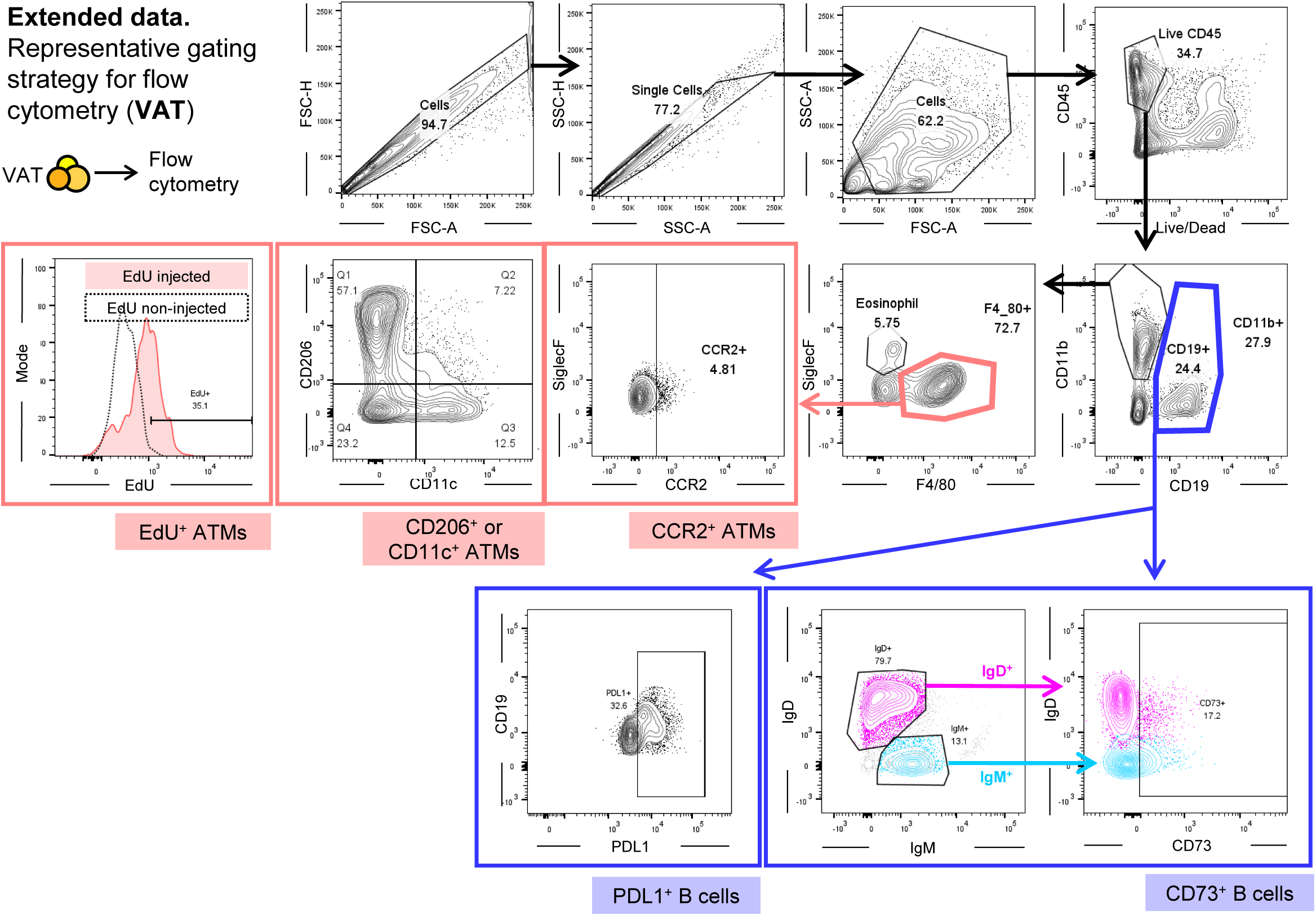

